# Phase-space distance between stationary states modulates phenotypic plasticity in breast cancer

**DOI:** 10.64898/2026.03.06.710190

**Authors:** Mayara D. A. Caldas, Alexandre A. B. Lima, Francisco Lopes

## Abstract

Phenotypic plasticity in breast cancer involves stochastic transitions between gene-expression states associated with clinically distinct subtypes, such as HER2+ and triple-negative breast cancer (TNBC). In non-conservative gene regulatory networks, however, the dynamical features controlling transition probabilities and timescales cannot be consistently reduced to potential-barrier depth, since potential functions are generally non-unique outside equilibrium. Here, by analyzing an ***NF-κB***-centered regulatory network that supports HER2+ and TNBC attractors, we identify a geometric control principle: the phase-space distance between stationary states, together with the bifurcation structure organizing multistability, provides a robust determinant of transition probabilities, times, and variability. We show that multistability is necessary but not sufficient for transitions, whose accessibility is constrained by the global arrangement of basins and the intervening unstable state. Within this framework, we find a marked asymmetry in dynamical sensitivity: the HER2+ regime is robust to intrinsic parameter variations, whereas the TNBC regime strongly amplifies such variations, offering a dynamical explanation for the pronounced gene molecular and clinical variability observed in TNBC. Together, our results establish a general geometric perspective on phenotypic transitions in non-conservative regulatory networks.

## 1 Introduction

Breast cancer is the most commonly diagnosed cancer worldwide and remains a leading cause of cancer-related mortality among women Sung et al. (2021). A central challenge in its clinical management is heterogeneity, both across patients and within individual tumors, which contributes to variability in prognosis, therapeutic response, and clinical outcomes Lüönd et al. (2021). In routine practice, tumors are commonly assigned intrinsic-like subtypes using immunohistochemical (IHC) surrogates for estrogen receptor (ER), progesterone receptor (PR), and human epidermal growth factor receptor 2 (HER2), together with the proliferation marker Ki-67, whereas molecular classification relies on multigene assays (e.g., PAM50) Maisonneuve et al. (2014). Among clinically defined categories, HER2-enriched tumors (HER2+) and triple-negative breast cancer (TNBC) represent contrasting phenotypic regimes, with TNBC lacking ER, PR, and HER2 expression and exhibiting particularly aggressive behavior and poor prognosis. Importantly, TNBC is also marked by pronounced molecular and clinical variability, motivating multiple subclassification schemes and underscoring the need for dynamical models that can capture its plasticity and unpredictability Lehmann et al. (2016); Burstein et al. (2015).

Recent single-cell RNA sequencing (scRNA-seq) studies have further sharpened this challenge by revealing extensive intra-tumoral heterogeneity at the transcriptomic level Chung et al. (2017); Wu et al. (2021). In particular, clinically classified tumors can contain subpopulations whose expression profiles resemble other subtypes, suggesting that tumor cell populations may occupy a repertoire of phenotypic states rather than a single homogeneous identity. Such observations motivate a dynamical perspective in which phenotypes correspond to stable gene-expression states generated by underlying regulatory circuitry, and heterogeneity reflects stochastic fluctuations within and between their basins of attraction. In this view, an important mechanistic question is not only whether multiple stable states coexist, but which dynamical features determine the accessibility, probability, and timescales of transitions between them.

The cancer attractors paradigm formalizes this idea by representing phenotypic states as attractor basins in an epigenetic landscape Waddington (1957); Kauffman (1971); Huang et al. (2009). Within this metaphor, transitions are commonly interpreted as barrier-crossing events governed by an effective potential, so that deeper basins or higher barriers imply lower transition probability and longer escape times. While conceptually powerful, this interpretation becomes problematic for gene regulatory networks, which operate far from equilibrium due to continuous synthesis and degradation, external driving, and dissipation. In such non-conservative systems, potential-like constructions are generally non-unique and do not uniquely determine the dynamics, limiting the use of barrier height as a universal control variable for transition probabilities. This motivates alternative descriptors of transition propensity that remain well-defined in non-equilibrium regulatory networks and can be directly related to model parameters.

Nuclear factor kappa B (*NF*-*κB*) provides a compelling biological context to examine these questions. *NF*-*κB* is a family of transcription factors broadly expressed across cell types and centrally involved in inflammatory and stress-responsive gene expression programs. In breast cancer, constitutive or dysregulated *NF*-*κB* activity has been associated with aggressive disease and therapy resistance, and has been implicated in processes such as epithelial-to-mesenchymal transition (EMT), stem-like features, and remodeling of the tumor microenvironment Pavitra et al. (2023); Wang et al. (2015). These observations have motivated mechanistic models of NF-*κ*B regulation aimed at connecting molecular circuitry to emergent phenotypic behavior. Based on experimental evidence and well documented regulatory interactions Pires et al. (2017); Sung et al. (2014); Pahl (1999), we previously developed a gene regulatory network (GRN) model in which NF-*κ*B participates in its own regulation together with key EMT-related genes Lopes et al. (2025). This model exhibits two stable fixed points corresponding to gene-expression profiles associated with HER2+ and TNBC phenotypes, separated by an unstable steady state and surrounded by distinct basins of attraction. Importantly, stochastic fluctuations in gene expression can drive spontaneous, irreversible transitions from the HER2+ attractor to the TNBC attractor at randomly distributed times, providing a mechanistic route to intra-tumoral heterogeneity.

Here, we build on this framework to systematically investigate what controls transition probabilities and timescales under parameter variation, with an emphasis on quantities that remain meaningful in non-conservative settings. To isolate the dynamical core responsible for multistability, we focus on a simplified sub-model containing only the *NF*-*κB* self-regulatory mechanism, whose positive feedback constitutes the primary driver of bistability and switching. This reduction enables a comprehensive analysis of how kinetic parameters shape (i) the existence and geometry of the bistable regime and (ii) the accessibility of stochastic transitions between basins.

We first perform single-parameter bifurcation analyses to map stationary states and identify bistable regions, while quantifying how the HER2+ and TNBC branches respond to intrinsic parameter perturbations. We then connect deterministic stationary-state geometry to cellular-scale switching behavior using stochastic simulations, and evaluate how transition probabilities, times, and variability depend on the separation between stable and unstable stationary states in phase space. Finally, we extend the analysis to two-parameter bifurcation diagrams to characterize how combined perturbations reshape the global organization of multistability, including qualitative differences in cusp geometry that delimit the parameter regions compatible with phenotypic plasticity.

Our results support a geometric control principle for phenotypic transitions in non-conservative regulatory networks. Specifically, we show that the coexistence of two stable states is only a necessary, but not sufficient, condition for transitions: accessibility depends on the global arrangement of basins and on the phase-space distance between stable states and the intervening unstable state. Within this geometric framework, we find a marked asymmetry in dynamical sensitivity: across parameters, the HER2+ branch is comparatively robust, whereas the TNBC branch strongly amplifies intrinsic parameter variability, offering a dynamical interpretation for the pronounced molecular and clinical variability observed clinically and experimentally in TNBC Lehmann et al. (2016); Burstein et al. (2015). More broadly, by replacing potential-barrier intuitions with distances and bifurcation geometry, quantities that remain well-defined far from equilibrium, our analysis provides a general route to quantify transition propensity and phenotypic plasticity in gene regulatory networks.

## 2 Results

### 2.1 Breast cancer subtypes map onto two basins of attraction

We study a gene regulatory network model describing the self-regulation of *NF*-*κB*, originally introduced in Lopes et al. (2025). The model focuses on the role of the transcription factor *NF*-*κB* in driving breast cancer heterogeneity. Specifically, it describes the regulation of the *p*50 and *p*65 genes by *NF*-*κB*, which in turn encode the synthesis of the *p*50 and *p*65 subunits of *NF*-*κB* itself. By capturing this positive feedback loop, the model incorporates two mechanisms that have been shown to play critical roles in gene regulation and development: positive feedback Lopes et al. (2008) and dimer formation Hsu et al. (2016). This framework allows us to investigate how *NF*-*κB* dynamics contribute to cancer cell progression and phenotypic heterogeneity.

Figure 1 depicts the gene regulatory network. Its components and reactions are described below:

- *N*0*X*: Denotes the unbound (unoccupied) regulatory region of gene *X*.
- *N*1*X*: Denotes the region of gene *X* bound by an *NF*-*κB* dimer.
- Gene *X*: *p*50 or *p*65.

Gene activation is described by a three-step process:

- Reversible binding of *NF*-*κB* to the gene’s regulatory region: *NF*-*κB* + *N*0*X* ↔ *N*1*X*.
- Irreversible RNA synthesis: *N*1*X* → *N*1*X* + *RNAX*.
- Irreversible protein synthesis: *RNAX* → *RNAX* + *X*.

In addition, the model describes the following constitutive reactions:

- Constitutive RNA synthesis: 0 → *RNAX*.
- RNA degradation: *RNAX* → 0.
- Protein degradation: *X* → 0.
- Dimer formation: *NF*-*κB* ↔ *p*50 + *p*65.

**Fig. 1:**
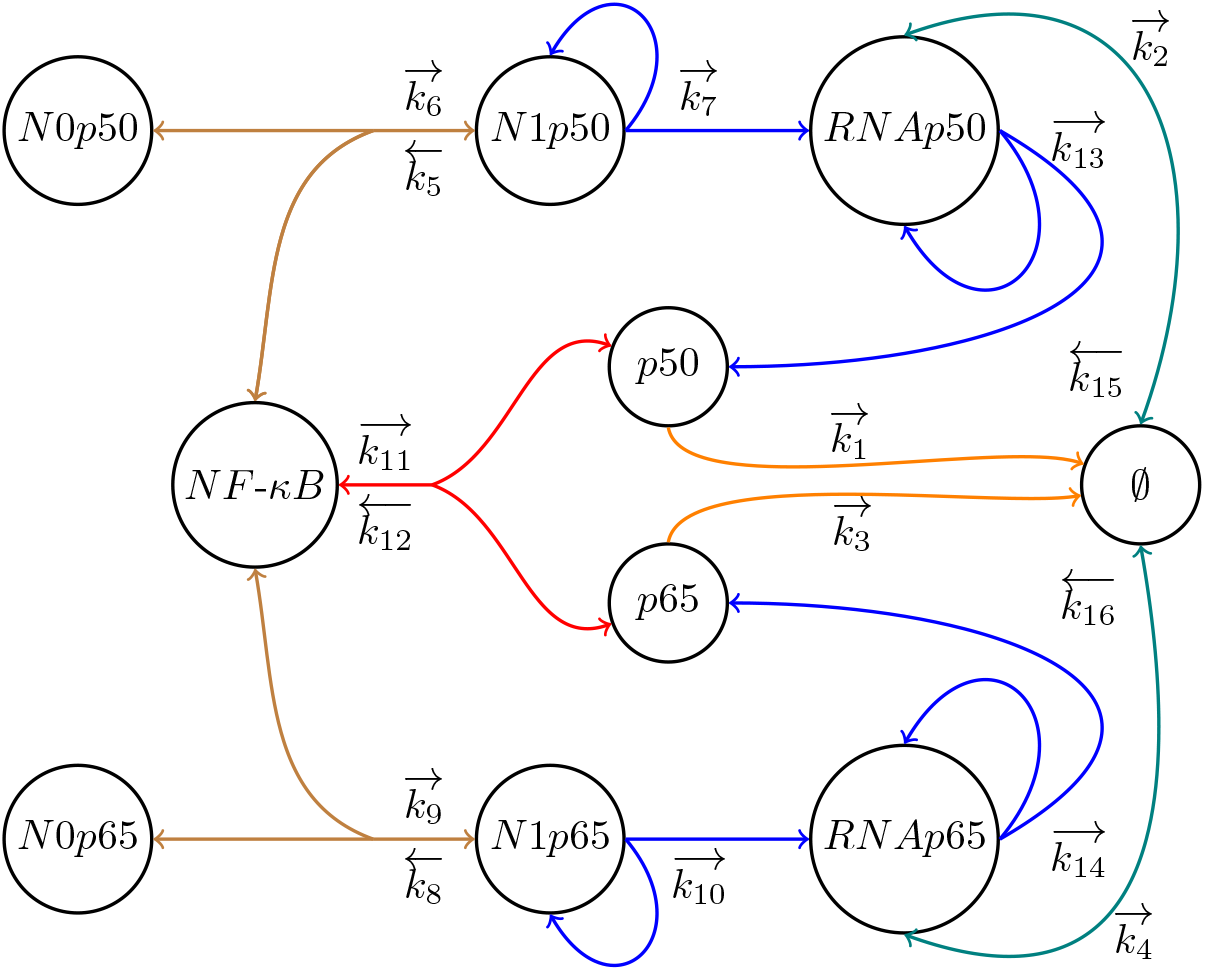
*NF*-*κB* Gene Regulatory Network. Brown lines represent *NF*-*κB* binding to the *p*50/*p*65 gene promoter. Red represents dimerization. Blue indicates RNA and protein synthesis. Orange represents degradation. Teal indicates constitutive synthesis and degradation. The kinetic constant 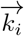 and 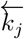 are associated to the forward and reverse reactions respectively. See Table S1 for traditional representation.

These sequential reactions describe the transcription factor *NF*-*κB* binding to the promoter region of the *p*65 gene, promoting its transcriptional activation. Once activated, the gene is transcribed into messenger RNA, which is subsequently translated into the *p*65 protein. A similar process occurs with the *p*50 gene: *NF*-*κB* binds to its promoter, inducing the production of *RNAp*50 and consequently the corresponding *p*50 protein. These two proteins, *p*50 and *p*65, associate in the cytoplasm to form new *NF*-*κB* dimers. This establishes a positive feedback loop in which *NF*-*κB* activates the expression of the very genes that encode its own subunits, thereby reinforcing its activity and enabling sustained regulatory dynamics.

By applying the law of mass action (Espenson (1995)) to the reactions described in the regulatory network (Figure 1) we obtain the following system of ordinary differential equations:

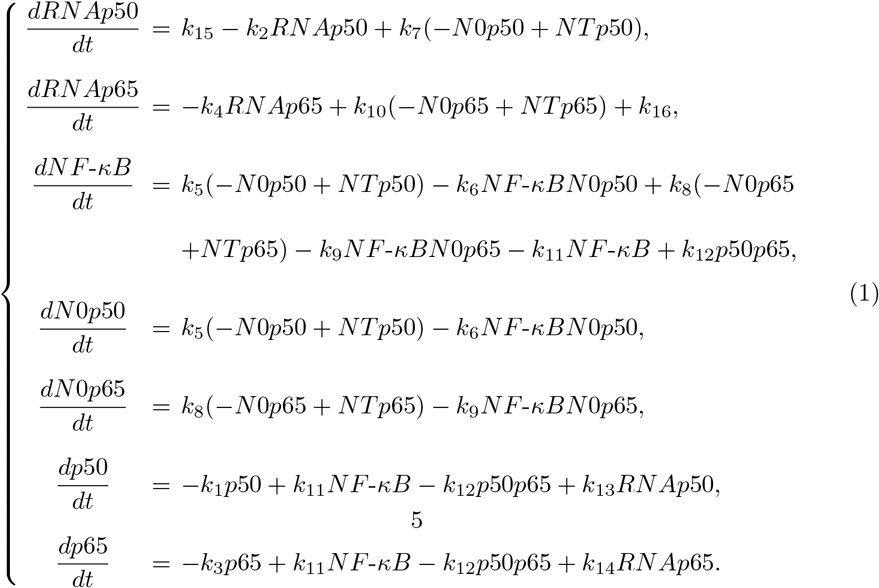

From Lopes et al. (2025), we used

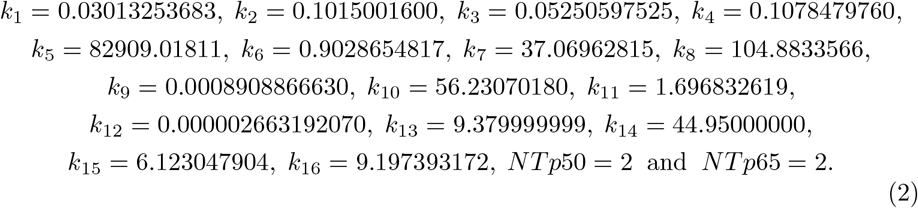

All kinetic parameters are treated as dimensionless in the main text for simplicity, with their corresponding dimensional forms and units reported in Table S1. The model admits three stationary points. A linear stability analysis shows that two are locally asymptotically stable, with all eigenvalues having negative real parts, and one is unstable, with one eigenvalue having a positive real part. This configuration characterizes the system as bistable, since it possesses two stable steady states separated by an unstable one. The parameters in (2) were determined using experimental data on the RNA levels of *NF*-*κB* in two well-known cell lines, HCC-1954 and MDA-MB-231, classified as subtypes HER2+ and triple negative breast cancer (TNBC), respectively. A key characteristic of these cells is the high level of *NF*-*κB* in the TNBC subtype compared to HER2+. The model bistable behavior was key for the model calibration because it allowed for the same set of parameter to reproduce both stable levels of *NF*-*κB* identified in each BC subtype cell lines. Both stable states are represented in Figure 2, together with the unstable one. Below, we present the stationary points, listing the copy number of *RNAp*50, *RNAp*65, *NF*-*κB, N* 0*p*50, *N*0*p*65, *p*50, and *p*65, respectively:

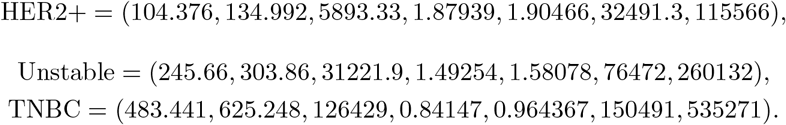

**Fig. 2:**
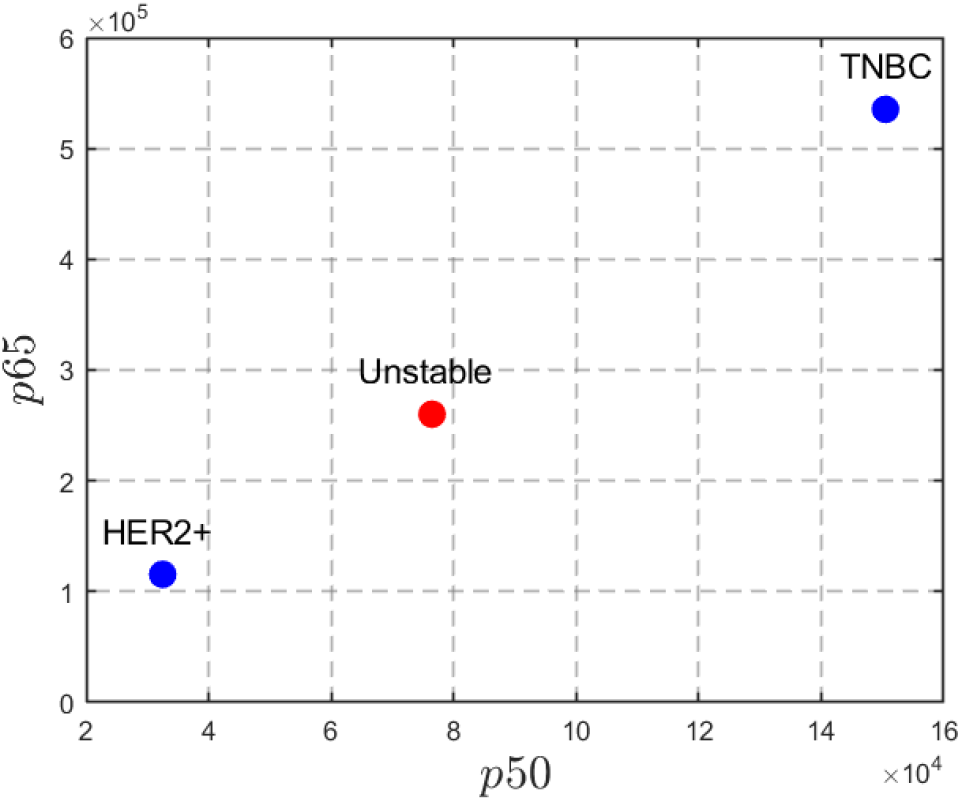
Projection of the system’s stationary states onto the *p*50 × *p*65 plane, highlighting the coexistence of two stable phenotypic states (HER2+ and TNBC) separated by an unstable saddle point. Expression levels are shown as absolute numbers of molecule copies.

The ability of our bistable model to reproduce *NF*-*κB* expression profiles of two breast cancer subtypes provides evidence for the cancer attractor framework, in which cellular phenotypes correspond to stable attractor states shaped by gene regulatory dynamics.

Considering the calibrated model, we computed the Euclidean distance between the stationary states in the phase space (i.e., the shortest straight-line distance between two points in that space), where each axis corresponds to the absolute molecule count of a species. This distance, reported in molecule copy number (c.n.), quantifies the aggregate change in molecular abundance required to traverse between states. The distance between the HER2+ state and the unstable point is 153216.35 m.u., while for the TNBC state it is 300407.21 m.u., indicating that the unstable point is much closer to the HER2+ state, almost half the distance compared to TNBC. Notably, stochastic single-cell simulations by Lopes et al. (2025) demonstrated that HER2+ tumors can evolve into the TNBC subtype, with 120 simulated trajectories showing this transition over a 20-year period.

These findings raise an important question: how can parameter variation increase these distances and potentially prevent transitions between subtypes?

To address this, in the next section we show bifurcation analyses with respect to each system parameter, enabling us to evaluate how the distances between the stationary states depend on parameter values.

### 2.2 Bifurcation analysis shows that TNBC is highly sensitive to parameter variation

Our bifurcation analysis aimed to map how the parameter space governs phenotypic stability. To this end, we first characterized how variations in key parameters modulate the concentrations of species within each basin of attraction, thereby assessing the sensitivity of each attractor to perturbations. This analysis was performed using the numerical continuation software MatCont (A. Dhooge and W. Govaerts and Yu.A. Kuznetsov and H.G.E. Meijer and B. Sautois (2003–2023)), a numerical tool widely used for characterizing bifurcations in systems of ordinary differential equations. Initially, we examined the effect of varying the parameter *k*_3_, which corresponds to the degradation reaction *p*65 → 0, and its influence on the dynamics of *NF*-*κB*.

The continuation analysis reveals that saddle-node bifurcations control the phenotypic transition between HER2+ and TNBC breast cancer subtypes, with the parameter *k*_3_ playing a critical role in this process by modulating *NF*-*κB* expression levels. Figure 3 displays the *NF*-*κB* bifurcation diagram with respect to *k*_3_, high-lighting regions of monostability and bistability. This diagram exhibits the inverted S-shaped curve, characterized by two saddle-node bifurcations that delimit three distinct regions of the parameter *k*_3_:

- For *k*_3_ ∈ (0.0403, 0.048), the system has a monostable region, associated with the TNBC phenotype.
- For *k*_3_ ∈ (0.048, 0.056), the system exhibits bistability, characterized by the coexistence of two stable solutions (HER2+ and TNBC states) separated by an unstable point.
- For *k*_3_ ∈ (0.056, 0.0655), the system exhibits only a monostable region, in this case associated with the HER2+ phenotype.

**Fig. 3:**
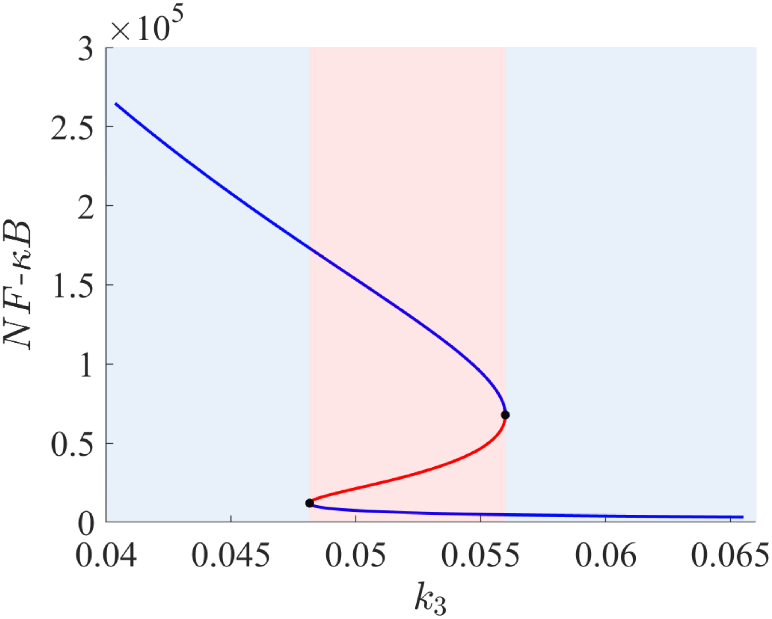
Bifurcation diagram of *NF*-*κB* with respect to *k*_3_. Stable solutions are represented in blue, while unstable solutions are represented in red.The red region represents bistability and the blue regions represent monostability.

The geometry of the bifurcation curve reflects different sensitivities associated with the TNBC and HER2+ phenotypes. Its lower branch, corresponding to the HER2+ phenotype, is characterized by low levels of *NF*-*κB* and by a relatively small variation in its concentration across a wide range of *k*_3_ values. This behavior indicates that, in this regime, the system is only weakly sensitive to changes in the degradation rate of *p*65. As a result, small variations in *k*_3_ are not sufficient to significantly perturb the *NF*-*κB* levels. In contrast, the upper branch, associated with the TNBC phenotype, exhibits a steeper slope, reflecting a much higher sensitivity to changes in *k*_3_. Thus, variation in *k*_3_, which changes the degradation rate of *p*65, leads to a marked variation in the stationary level of *NF*-*κB*.

Figure 4 shows the bifurcation diagrams for all remaining variables with respect to *k*_3_. The variables *RNAp*50, *RNAp*65, *p*50 and *p*65 all exhibit the same inverted S-shaped curve as *NF*-*κB*. This trend occurs because *k*_3_ sets the degradation rate of *p*65: higher values of *k*_3_ lead to lower steady?state levels of these variables, which is characteristic of the HER2+ subtype. In contrast, the diagrams for *N*0*p*50 and *N*0*p*65 display the opposite pattern, that is, an S-shaped curve. This behavior occurs because *N*0*p*50 and *N*0*p*65 represent unoccupied gene regulatory regions; therefore, a greater *k*_3_ value leads to a lower amount of *NF*-*κB*, making more of these regions available (i.e., unbound).

**Fig. 4:**
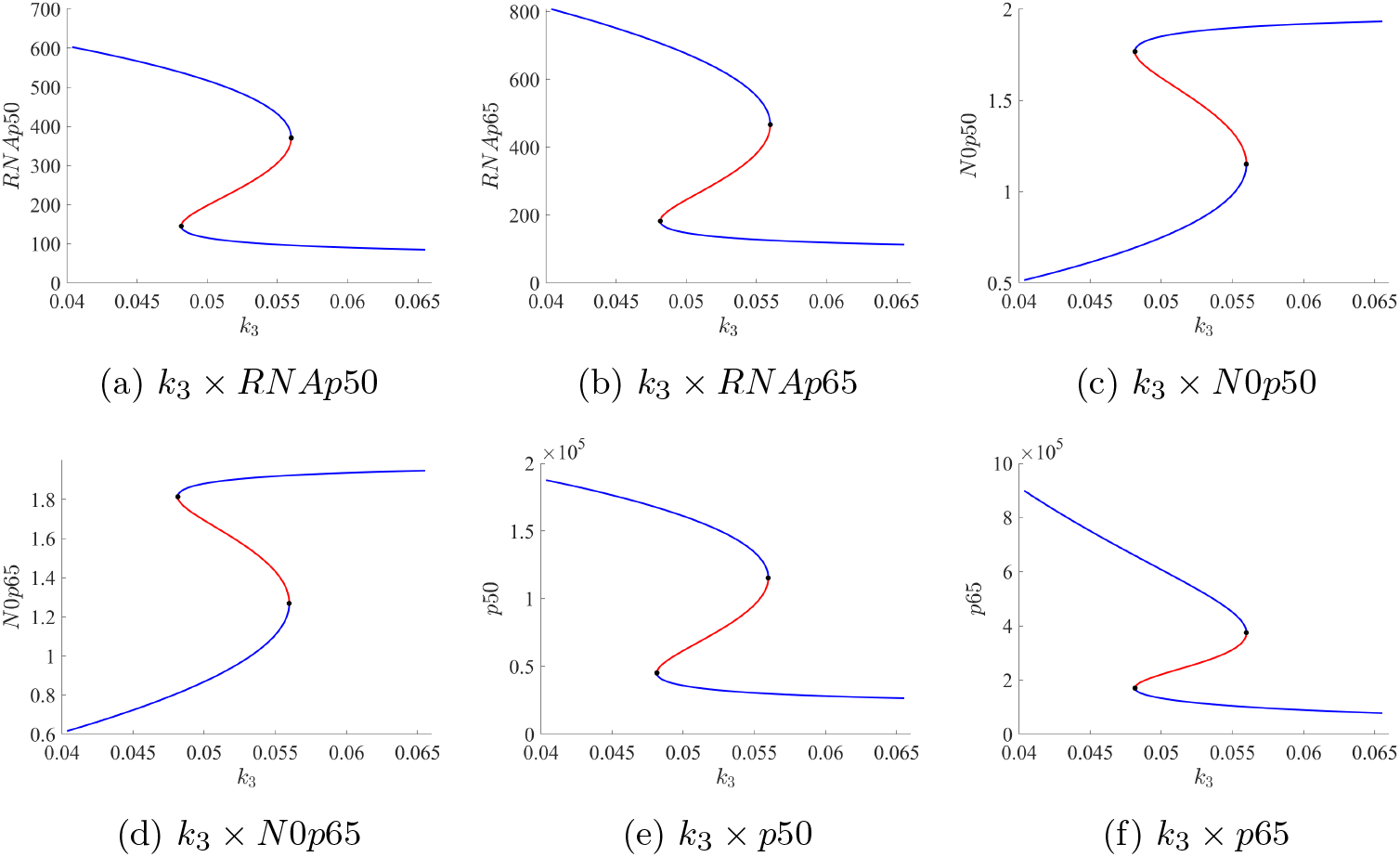
Bifurcation diagrams of all species, except *NF*-*κB*, with respect to *k*_3_. Stable solutions are represented in blue, while unstable solutions are represented in red.

To determine how the system’s variables are affected by the parameter *k*_3_ along the different bifurcation branches, we performed a sensitivity analysis. The results are summarized in Table 1. We also show the percentage variation for each branch relative to the total variation (set to 100%). Figure 5 shows the relative sensitivity of each variable to *k*_3_, indicating that, in the TNBC branch, all variables are highly sensitive to this parameter.

**Table 1:** Variation of each species’ copy number along the HER2+ and TNBC branches as a function of *k*_3_ in the corresponding bifurcation diagrams. The columns are as follows: (1) variables; (2) absolute variation along the TNBC branch; (3) absolute variation along the HER2+ branch; (4) total absolute variation across the entire bifurcation diagram; (5-6) the percentage values, with column (4) representing 100%.

| Variables | TNBC | HER2+ | Total | % TNBC | % HER2+ |
| --- | --- | --- | --- | --- | --- |
| $RN Ap50$ | 231.99 | 60.27 | 517.52 | 44.83 | 11.65 |
| $RN Ap65$ | 340.60 | 69.40 | 693.96 | 49.08 | 10.00 |
| $NF-\kappa B$ | 196895.13 | 8877.67 | 261499.54 | 75.29 | 3.39 |
| $N0p50$ | 0.64 | 0.16 | 1.42 | 44.83 | 11.65 |
| $N0p65$ | 0.65 | 0.13 | 1.33 | 49.08 | 10.00 |
| $p50$ | 72216.78 | 18762.52 | 161098.68 | 44.83 | 11.65 |
| $p65$ | 524527.20 | 92805.19 | 821504.67 | 63.85 | 11.30 |

**Fig. 5:**
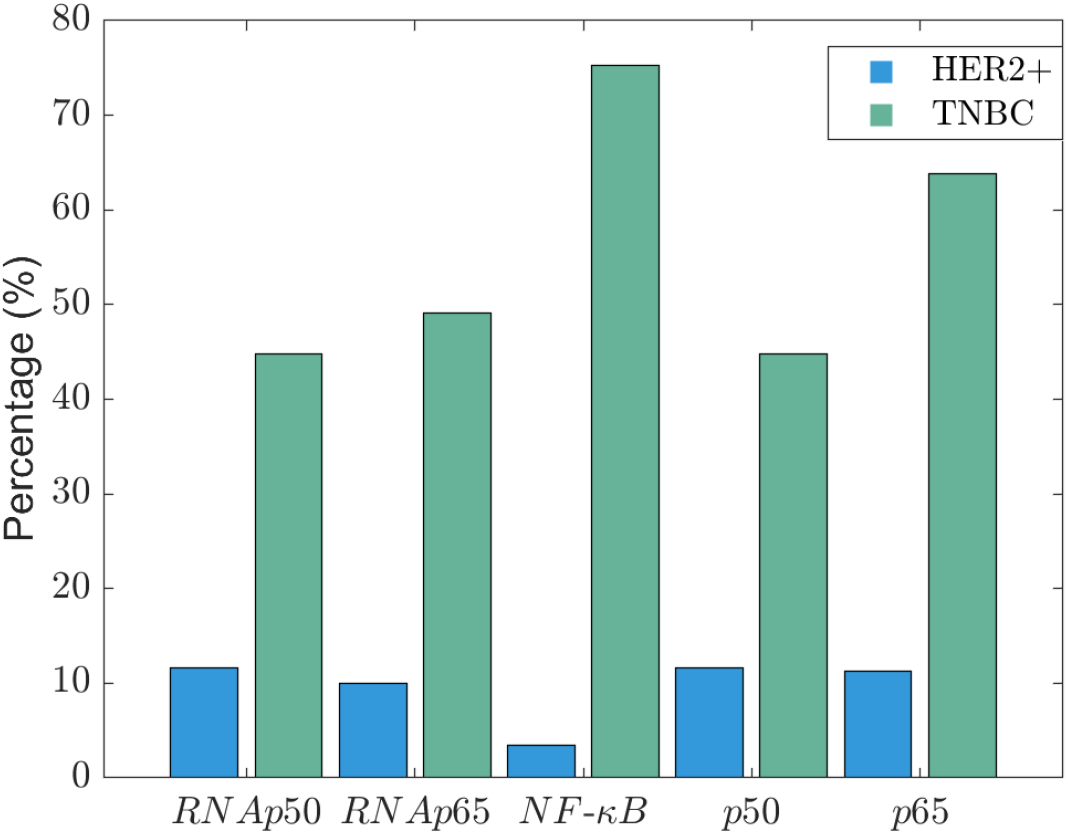
Sensitivity of each species to parameter *k*_3_. Blue and green bars represent the sensitivity based on the total percentage variation in the HER2+ and TNBC subtype branches, respectively.

Interestingly, although *p*65 would be expected to be more sensitive to *k*_3_ since this parameter is related to its degradation rate, *NF*-*κB* instead shows higher sensitivity in the TNBC branch. This unexpected behavior stems from the combined effect of dimerization to form *NF*-*κB* and its role in *p*50 and *p*65 synthesis. A reduction in *p*65 causes a reduction in the levels of *NF*-*κB* dimers, which results in lower levels of *p*50 and *p*65. Consequently, with fewer monomers available, the level of *NF*-*κB* is doubly affected. This high sensitivity may be linked to a clinical characteristic of TNBC: its significant variability among breast cancer subtypes. Consequently, several new TNBC subclassifications have recently been proposed. We explore this potential connection further in the Discussion.

In contrast to the sensitivity pattern in the TNBC branch, the concentration of all species in the lower branch, associated with HER2+, exhibits remarkably lower sensitivity to *k*_3_ (compare the percentage variation for each variable, Table 1). This indicates that gene expression levels in this subtype are more stable than in TNBC. The difference is even more pronounced for *NF*-*κB* concentration: the TNBC branch exhibits a 75.29% variation, compared to only 3.39% for the HER2+ branch. This suggests that, in the low protein level regime, the combined effects of dimerization and *NF*-*κB* play no significant role. This has a strong downstream effect, indicating that the expression levels of *NF*-*κB* target genes are expected to show lower sensitivity to *k*_3_, even when compared to the sensitivity of *p*50 and *p*65 levels.

The analysis for the parameter *k*_3_ can be extended to the remaining parameters in the GRN, Figure 1. Bistable and monostable intervals of all parameters are provided in Table S2, within the “Bifurcation diagrams with a free parameter” section of the Supplementary Material. The intervals of values for all variables on both stable branches, along with the respective bifurcation diagrams for each model parameter, are also presented (Tables *S*3, *S*4 and *S*(*N*+ 1) and Figures *S*1, *S*2 and *S*(*n* − 1) for *k*_1_, *k*_2_, and *k*_*n*_, *n* = 4, …, 16, respectively). The sensitivity analyses are shown in Figure S16. Selected cases from these results are discussed below.

All *NF*-*κB* bifurcation diagrams with respect to the kinetic parameter associated with the forward self-activation reactions, together with constitutive RNA synthesis, exhibit an S-shaped curve (see Figures *Si, i* = 5, 6, 8, 9, 11, 12, 13, 14, 15). This is expected since all these reactions lead to increasing *NF*-*κB* levels. Consequently, the higher these parameters are, the higher the *NF*-*κ*B levels will be, leading the upper side of the S-shaped curve to the right side. Conversely, the bifurcation diagram with respect to the kinetic parameter associate to all reactions contributing to reducing *NF*-*κB* levels show inverted S-shaped curves (Figure 3 and Figures *Si, i* = 1, 2, 3, 4, 7, 10). Interestingly, while the reactions *N*1*p*50 → *N*0*p*50 + *NF*-*κB* and *N*1*p*65 → *N*0*p*65 + *NF*-*κB*, associated with the kinetic parameters *k*_5_ and *k*_8_, respectively, initially appear to increase *NF*-*κB* levels, they ultimately reduce them because these reactions decrease the levels of *N*1*p*50 and *N*1*p*65. This leads to reduced synthesis of *p*50 and *p*65 and, consequently, lower *NF*-*κB* levels. Besides that, since the maximum amount of *N*1*p*65 and *N*1*p*50 is 2 (corresponding to both gene copies being bound), the contribution of these reactions towards increasing *NF*-*κB* is insignificant. In all cases, the upper branch corresponds to the TNBC subtype, whereas the lower branch corresponds to the HER2+ subtype. Moreover, in all the *NF*-*κB* diagrams, the lower branch is consistently less sensitive to parameter variations, whereas the upper branch shows greater responsiveness.

Figure S16 shows the sensitivity analysis for the bifurcation diagram of all species with respect to all parameters, except *k*_3_. It reveals that in the TNBC branch, *NF*-*κB* is the most sensitive variable to parameter variations across all diagrams. Conversely, in the HER2+ branch, it is the least sensitive variable in all diagrams. This indicates that in the HER2+ subtype branch, the variability of all variables is buffered at the level of *NF*-*κB*, while in the TNBC subtype branch, it is amplified. This pattern mirrors the one observed in the bifurcation diagrams with respect to *k*_3_ (Figure 5) and likely contributes to the high clinical variability found in TNBC-classified tumors.

### 2.3 The distance between phenotypic states modulates transition times

The arrangement between the stationary states, Subsection 2.1, reveal an important geometric feature: the unstable saddle point lies much closer to the HER2+ attractor than to the TNBC attractor, indicating an asymmetric separation between the two phenotypic basins. This observation raises the question of whether parameter variations could modify these distances and thereby prevent transitions between subtypes. To investigate this, we focused on parameters *k*_3_ and *k*_12_, which correspond to the reactions *p*65 → 0 and *p*50 + *p*65 → *NFκB*, respectively. These reactions have the strongest impact on *NFκB* levels. Since the analysis centers on distances between stationary points, it is restricted to the bistable region for each parameter. For various parameter values within the bistable region, we calculated the distances between the HER2+ and unstable point, as well as between the TNBC and unstable point. These distances were subsequently interpolated to produce continuous curves, Figure 6.

**Fig. 6:**
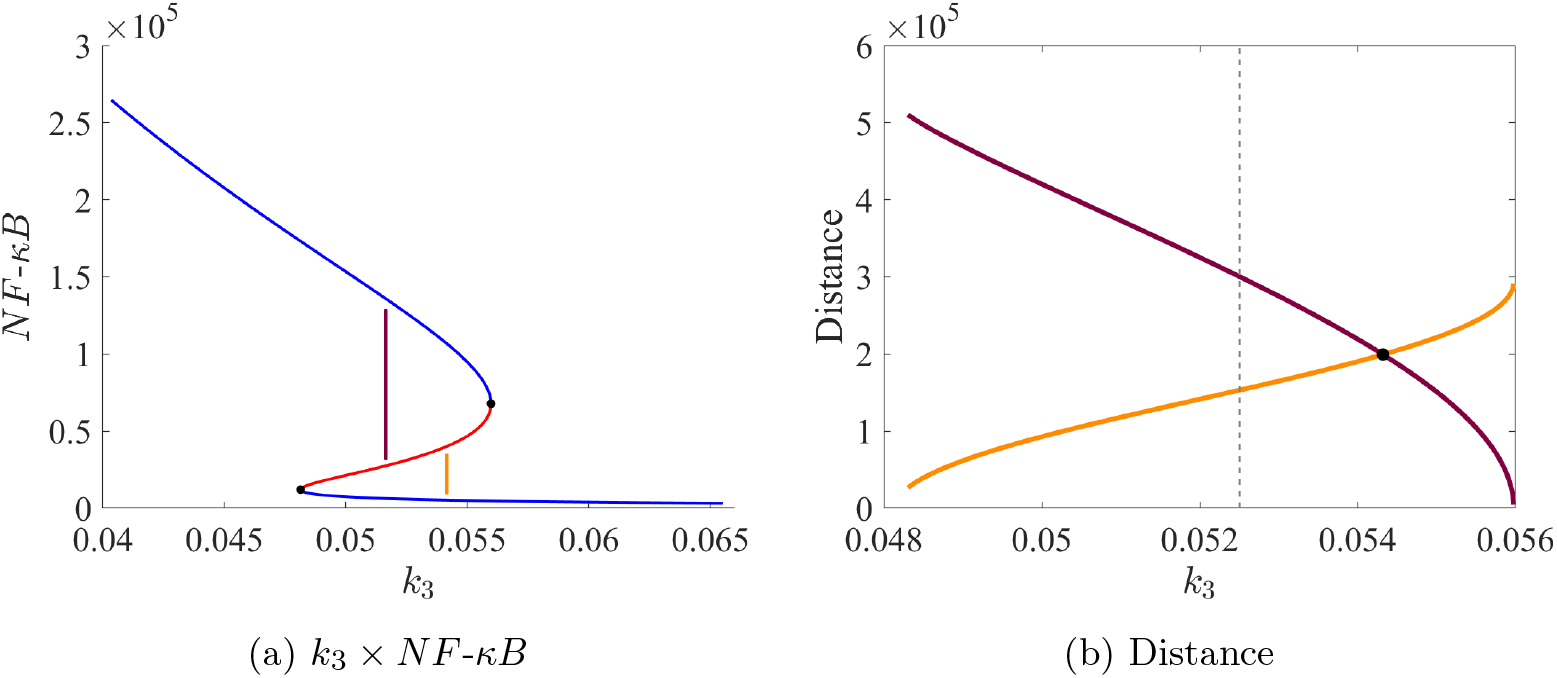
Variation of distances between stationary points with respect to *k*_3_. Brown and orange lines denote the distance from TNBC to the unstable point and from the unstable point to HER2+, respectively. (A) Bifurcation diagram of *k*_3_, indicating the distance between stationary states. (B) Variation of these distances, reported in c.n., as a function of *k*_3_, with the dashed line corresponding to the value of *k*_3_ in the original system. (*k*_3_ = 0.05250597525).

Figure 6a shows the *NF*-*κB* bifurcation diagram with respect to *k*_3_ indicating how the distance between states were determined. From the left border of the bistable region toward higher *k*_3_ values, the distance between the HER2+ and the unstable state increases gradually (Figure 6b, orange curve), while the distance between the unstable and the TNBC state decreases, brown curve. Both distances become identical (199405.21 c.n.) at *k*_3_ = 0.054321555749, with a relative error of only 8.099 × 10^−10^ (i.e., 8.10 × 10^−8^%). This *k*_3_ value corresponds to a 3.46% increase compared to the originally calibrated value (*k*_3_ = 0.05250597525), Table 2. Such an increase in *k*_3_ results in a 30.15% increase in the distance between HER2+ and the unstable state, and a 33.62% decrease in the distance between the unstable and TNBC states.

**Table 2:** Distances, in c.n., from HER2+ and TNBC to the unstable state for different *k*_3_ values, and their percentage variations relative to the original.

| $k_3$ | Variation (%) | HER2+ ↔ Unstable | Variation (%) | TNBC ↔ Unstable | Variation (%) |
| --- | --- | --- | --- | --- | --- |
| 0.05250597525 | Not apply | 153216.35 | Not apply | 300407.21 | Not apply |
| 0.054321555749 | ↑ 3.46% | 199405.21 | ↑ 30.15% | 199405.21 | ↓ 33.62% |
| 0.0511933259 | ↓ 2.5% | 122670.94 | ↓ 19.94% | 364024.90 | ↑ 21.18% |

To further explore the role of this kinetic parameter for the transition between states, we tested the effect of a 19.94% reduction in the distance from HER2+ to the unstable state, from the original 153216.35 c.n. to 122670.94 c.n.. This was achieved by a 2.5% decrease in *k*_3_ from the originally value. The distance from the unstable state to TNBC was increased by 21.18%.

To assess the impact of these altered distances between stable states on subtype transitions, we performed 120 stochastic simulations using COPASI (see Hoops et al. (2006)). For the values of *k*_3_ that placed the unstable state at the midpoint between HER2+ and TNBC, none of the 120 simulations exhibited a transition from HER2+ to TNBC, even after 20 years of simulated time. A randomly selected simulation (Figure 7) shows the system fluctuating around the HER2+ stable state. This behavior was expected, since for this *k*_3_ value the distance from HER2+ to the unstable point is increased by 30.15% compared to the original calibration.

**Fig. 7:**
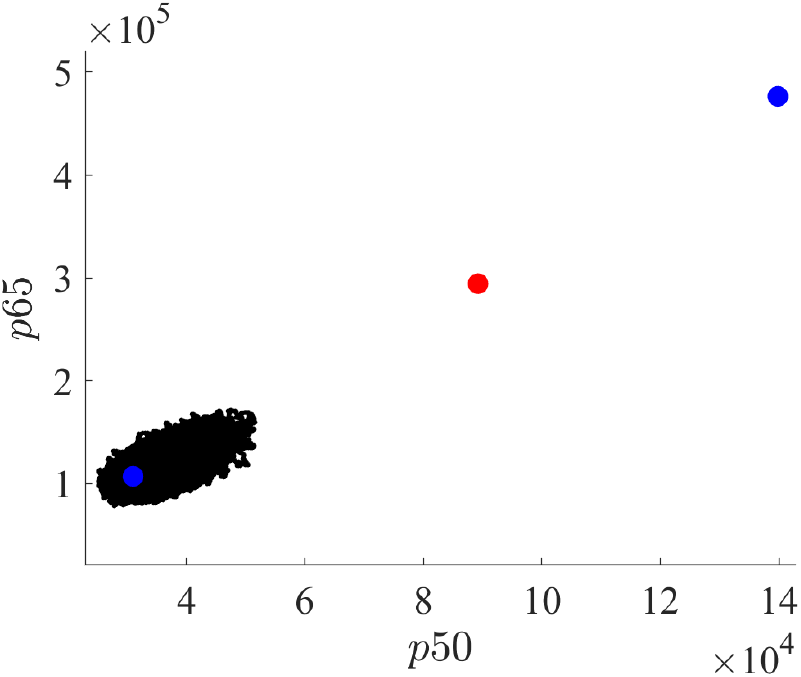
Stochastic model simulation for *k*_3_ = 0.054321555749, which places the unstable state at the midpoint between HER2+ and TNBC states. Over a time period of 20 years, the model remains fluctuating around HER2+.

For *k*_3_ = 0.0511933259 (a 2.5% reduction from the original value), which shortens the HER2+–unstable distance by 19.94%, all 120 simulations transitioned from HER2+ to TNBC within 18 days (Figure 8a). The trajectory in Figure 8b illustrates this rapid transition: the system spends only a brief time fluctuating around the HER2+ state before moving to the TNBC basin.

**Fig. 8:**
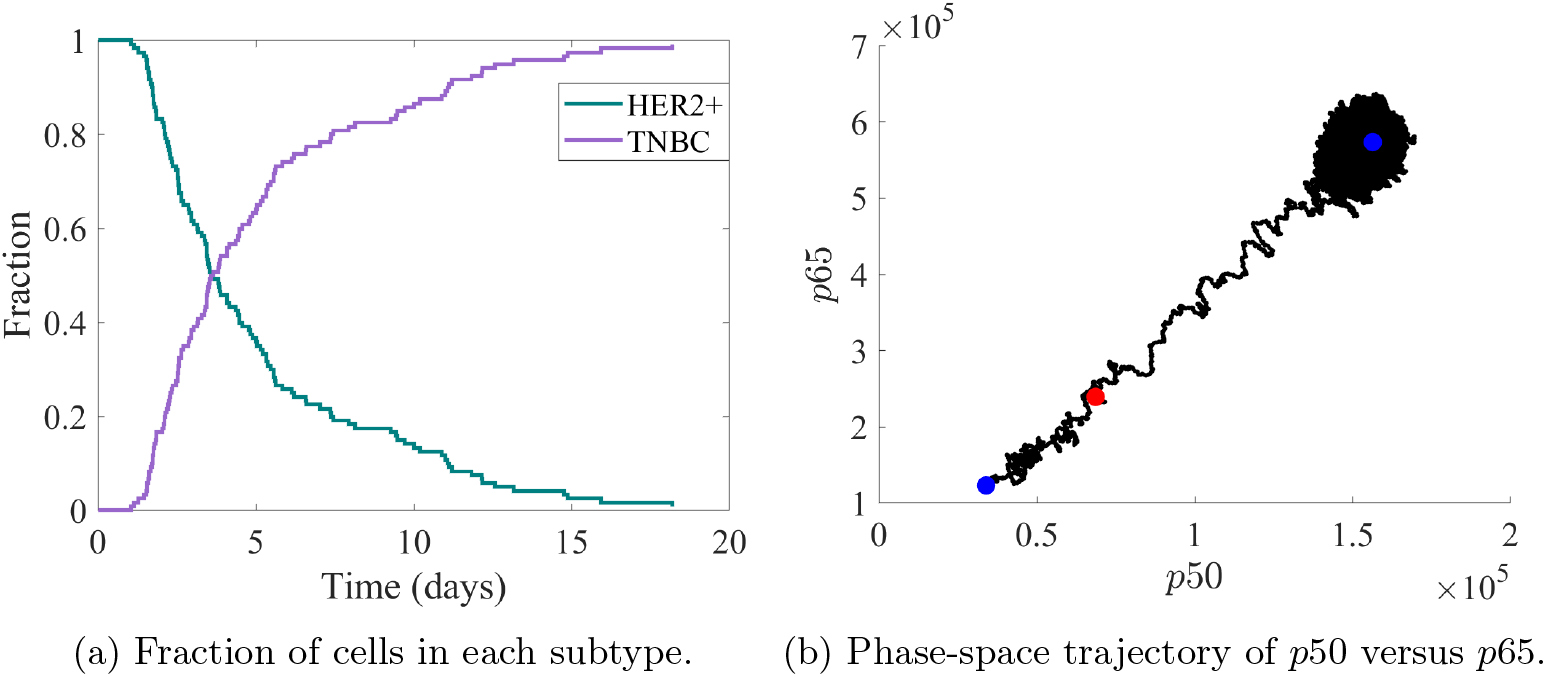
Stochastic simulations for *k*_3_ = 0.0511933259. (A) Fraction of HER2+ and TNBC cells over time across 120 simulations. All simulations start as HER2+ (fraction = 1) and complete the transition to TNBC (fraction = 1) within 18 days. (B) Transition trajectory from HER2+ to TNBC in the *p*50 versus *p*65 phase space.

In addition to *k*_3_, analyzed above, the dimer association parameter *k*_12_ is also critical for *NF*-*κB* levels (Figure 9a). However, unlike *k*_3_, for higher values of *k*_12_ the system settles into the TNBC attractor, exhibiting *NF*-*κB* levels characteristic of the TNBC subtype. Moving inside the bistable region from left to right increases the distance between TNBC and the unstable state (brown curve in Figure 9b). In this case, the value of *k*_12_ for which the unstable state is equidistant from HER2+ and TNBC is *k*_12_ = 2.574339 × 10^−6^, with a relative error of 4.5968 × 10^−6^ (i.e., 0.0004596814%), and the corresponding distance is 199405.21 c.n.. To verify the transition probabilities under these conditions, we performed 120 stochastic simulations with this value of *k*_12_. Over 20 years of simulated time, no transitions to the TNBC state were observed in any simulation, confirming the high stability of the HER2+ state. This behavior mirrors the stability observed when varying *k*_3_. As shown in the representative trajectory (Figure 10), the system fluctuates around the HER2+ steady state, reflecting the increased distance between it and the unstable state compared to the original configuration.

**Fig. 9:**
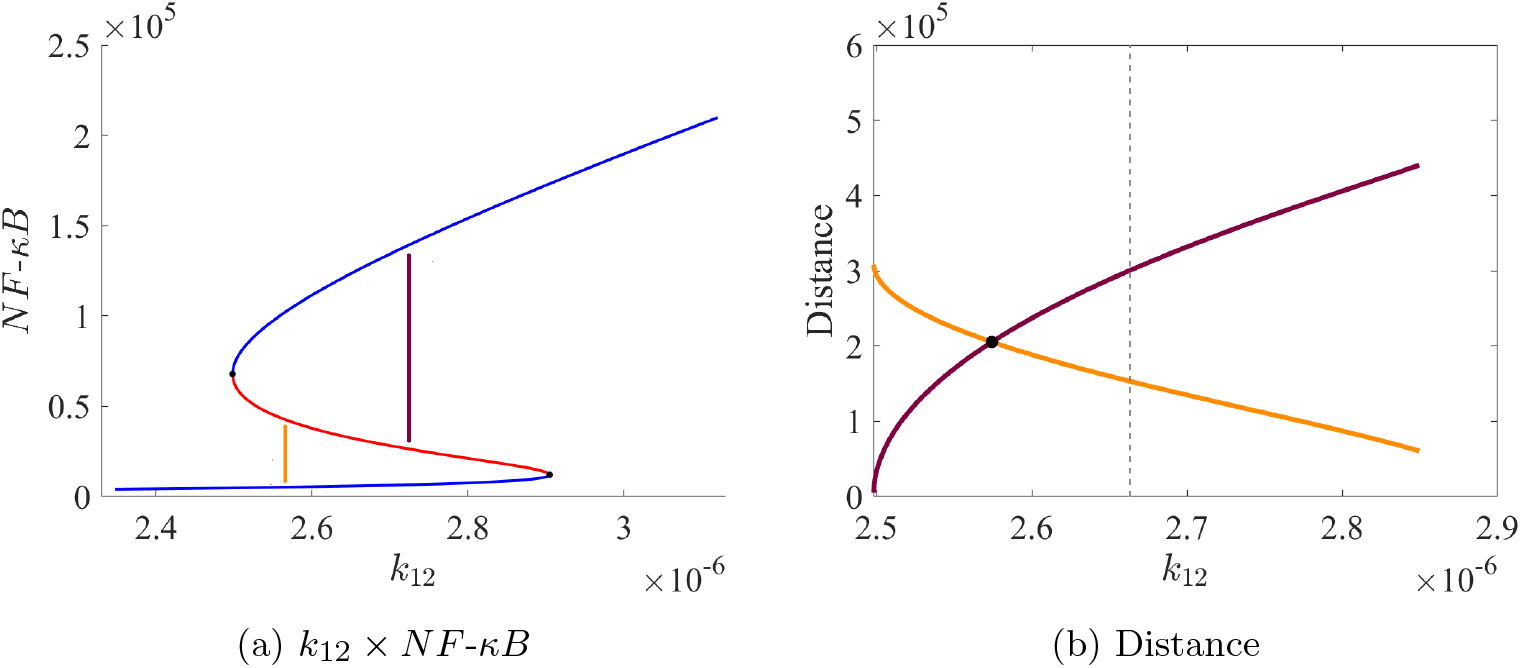
Distance with respect to *k*_12_. Brown and orange lines denote the distance from TNBC to the unstable point and from the unstable point to HER2+, respectively. Bifurcation diagram of *k*_12_, indicating the distance between stationary states. (B) Variation of these distances, reported in c.n., as a function of *k*_12_, with the dashed line corresponding to the value of *k*_12_ in the original system. (2.66319207 × 10^−6^).

**Fig. 10:**
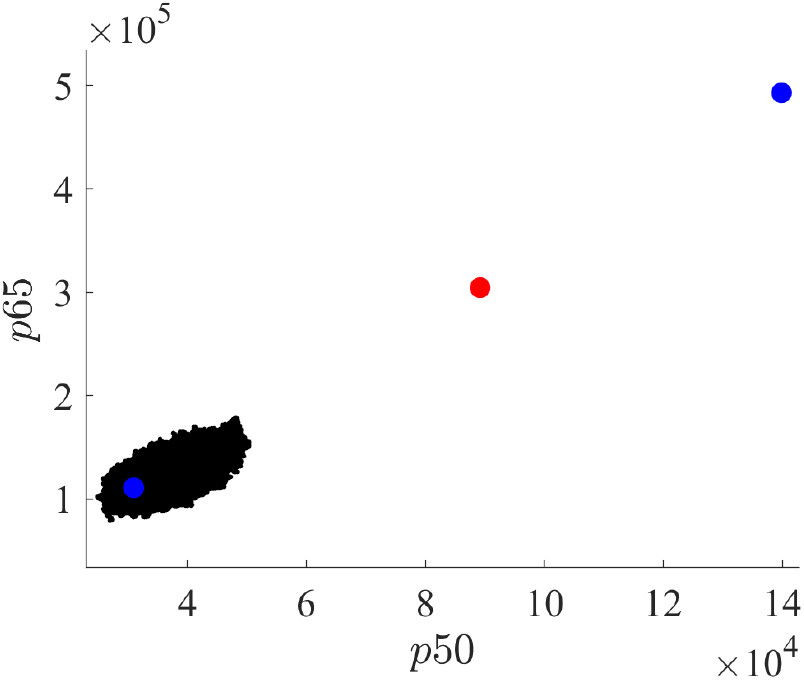
Representative stochastic simulation for the *k*_12_ value (2.574339 × 10^−6^) at which both HER2+-unstable and unstable-TNBC distances are equal (205386.83 c.n.). The simulation illustrates the persistence of the HER2+ state over 20 years.

It is remarkable how quickly the transition from HER2+ to TNBC occurs when the original HER2+?unstable state distance is reduced to 122670.94 c.n. by decreasing *k*_3_ by 2.5%; all 120 simulations transitioned in less than 18 days (Figure 8a). To investigate whether this behavior is preserved when the same distance is achieved by varying a different kinetic parameter, we increased the value of *k*_12_ by 2.35% such that the HER2+-unstable distance matched the same distance (122670.94 c.n.), Table 3. This reduced distance consistently facilitated rapid transitions, with all 120 simulations switching from HER2+ to TNBC within 32 days. As shown in the representative trajectory (Figure 11b), the system displayed only brief fluctuations around HER2+ state before transitioning to TNBC.

**Table 3:** Distances, in c.n., from HER2+ and TNBC to the unstable state for different *k*_12_ values, and their percentage variations relative to the original.

| $k_{12}$ | Variation (%) | HER2+ ↔ Unstable | Variation (%) | TNBC ↔ Unstable | Variation (%) |
| --- | --- | --- | --- | --- | --- |
| $2.66319207 \times 10^{-6}$ | Not apply | 153216.35 | Not apply | 300407.21 | Not apply |
| $2.574339 \times 10^{-6}$ | $\downarrow 3.33\%$ | 205386.83 | $\uparrow 34.05\%$ | 205385.89 | $\downarrow 31.63\%$ |
| $2.725683 \times 10^{-6}$ | $\uparrow 2.35\%$ | 122670.94 | $\downarrow 19.94\%$ | 351952.23 | $\uparrow 17.16\%$ |

**Fig. 11:**
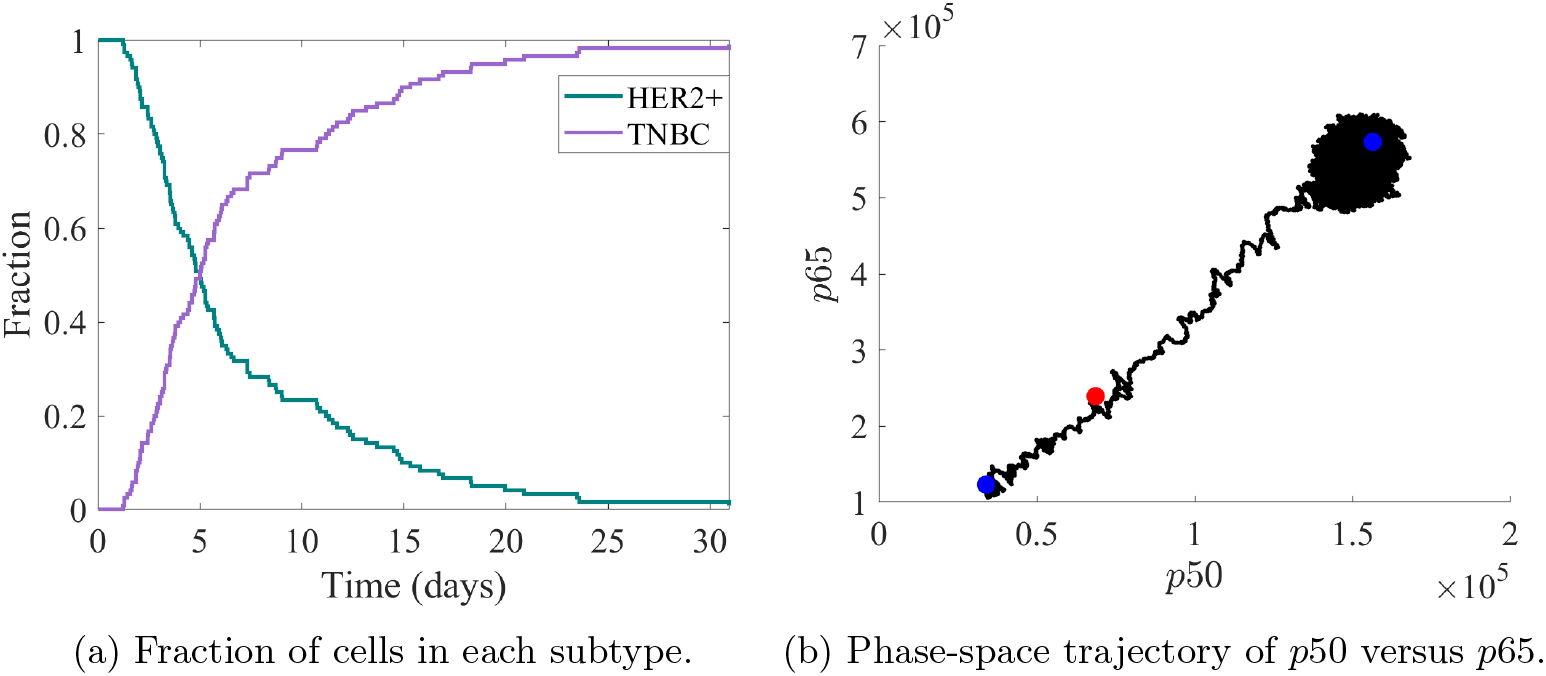
Stochastic simulation for a reduced distance between the HER2+ and unstable states (*k*_12_ = 2.725683 × 10^−6^). (A) Fraction of HER2+ and TNBC cells over time. All simulations start as HER2+ (fraction = 1) and complete the transition to TNBC (fraction = 1) within 32 days. (B) Phase-space transition trajectory from a randomly selected simulation.

Our analysis above showed that setting the HER2+-unstable distance to 122670.94 c.n., either by choosing *k*_3_ = 0.054321555749 or *k*_12_ = 2.725683 × 10^−6^, resulted in different transition times: 18 and 32 days, respectively. To deepen our analysis about these different behaviors according to the parameter changed, we fitted the distribution of the time of transition from each set of 120 simulations to log-normal probability density functions, as shown in Figure 12. The resulting statistical measures are summarized in Table 4.

**Table 4:** Statistical measures of transition times from 120 stochastic simulations for *k*_3_ and *k*_12_, both yielding the same HER2+-unstable state distance (122670.94 c.n.).

| Parameter | Mode (days) | Median (days) | Mean (days) | Variance (days<sup>2</sup>) |
| --- | --- | --- | --- | --- |
| $k_3$ | 2.416 | 3.885 | 4.928 | 14.772 |
| $k_{12}$ | 2.927 | 5.115 | 6.761 | 34.164 |

**Fig. 12:**
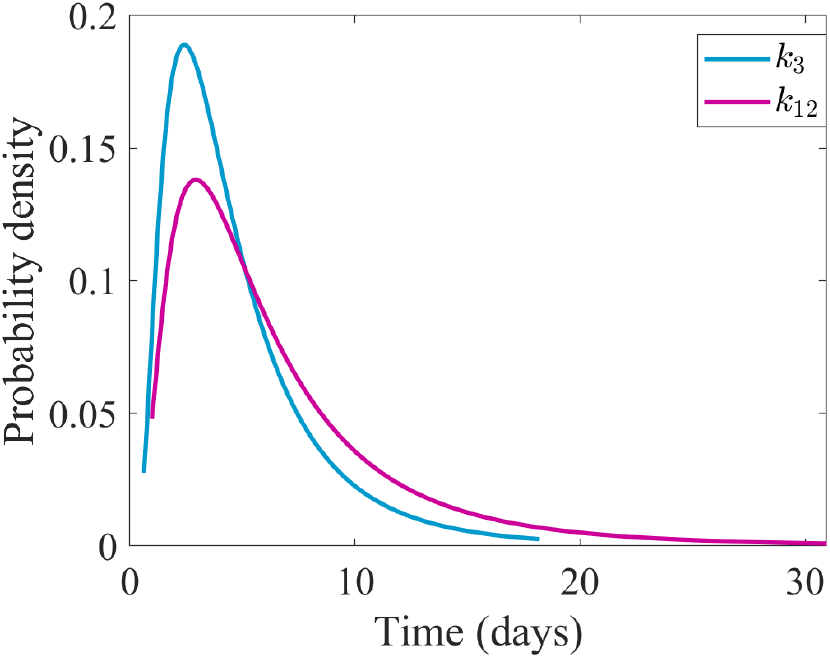
Log-normal distributions of transition times between the HER2+ and the unstable states, with the distance set to 122670.94 c.n.. Shown in teal-blue for *k*_3_ reduced to 0.0511933259 and in dark magenta for *k*_12_ increased to 2.725683 × 10^−6^.

The most probable transition time (mode) for *k*_3_ = 0.054321555749 is shorter than that for *k*_12_ = 2.725683 × 10^−6^, consistent with the full transition times (18 versus 32 days, respectively).

It is interesting to note that, to reach the 122,670.94 c.n. distance between the HER2+ and the unstable state, *k*_12_, associated with the reaction *p*50 + *p*65 → *NFκB*, was increased by 2.35%, whereas *k*_3_, associated with *p*65 → 0, was reduced by 2.5%. Together with the statistics presented in Table 4, this indicates that the network is more sensitive to variations in the reaction *p*50 + *p*65 → *NFκB* than in *p*65 degradation.

Notably, although variations in either parameter significantly accelerate transitions, changes in *k*_12_ lead to a broader distribution of transition times, reflecting increased variability in the system?s dynamics. In other words, perturbations in the dimerization reaction have a stronger and more heterogeneous impact on the transition process than changes in *p*65 degradation.

Taken together, these findings indicate that while the distance between stationary states is a critical determinant of transition probabilities, it is not the sole factor governing the system’s dynamics.

### 2.4 Parameter choice determines cusp geometry, resulting in unbounded or limited bistable regions

Following the previous single-parameter analysis, we extended the study to the simultaneous variation of two model parameters. This analysis enables the identification of regions in parameter space corresponding to distinct dynamic regimes and the characterization of bifurcation curves and critical boundaries. Together, these results provide a more comprehensive understanding of how pairs of parameters jointly shape the system’s global dynamics.

Since the levels of *NF*-*κB* and its subunits are critical for breast cancer progression, we first consider the simultaneous variation of parameters *k*_3_ and *k*_12_, which correspond to the degradation of *p*65 protein (*p*65 → 0) and the dimerization (*p*50 + *p*65 → *NF* - *κB*), respectively. The MatCont analysis (A. Dhooge and W. Govaerts and Yu.A. Kuznetsov and H.G.E. Meijer and B. Sautois (2003–2023)) reveals the presence of a cusp bifurcation: two saddle-node bifurcation lines delimiting a bistable region, whose origin corresponds to the cusp point (see Figure 13a). Within this region, the system exhibits bistability, whereas outside it the dynamics is monostable. The upper region outside the bistable region is associated to high levels of *NF*-*κB* corresponding to TNBC subtype, while the lower region corresponds to HER2+.

**Fig. 13:**
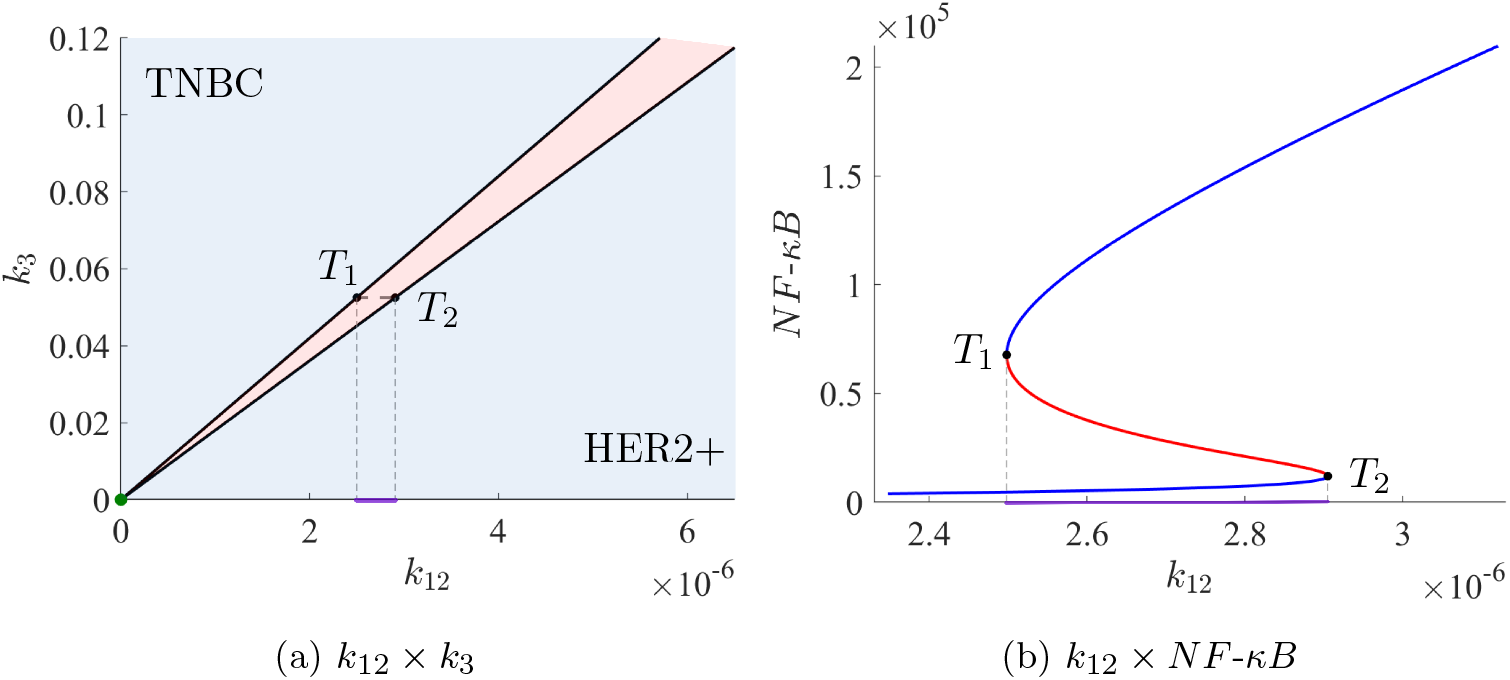
(A) Cusp bifurcation diagram with respect to *k*_3_ and *k*_12_. The green dot indicates the cusp point, the red region represents bistability, and the blue region represents monostability. The purple line corresponds to the limits of the bistable region for a given value of *k*_3_. (B) Bifurcation diagram with respect to *k*_12_ for *k*_3_ = 0.05250597525. The purple line corresponds to the same interval shown in (A).

It is worth noting that the bistable region in the cusp bifurcation diagram is an unbounded set. In particular, for any fixed *k*_3_ *>* 0, the bifurcation diagram with respect to *k*_12_, Figure 13b, exhibits the characteristic S-shaped curve, a typical signature of saddle-node bifurcations in bistable systems.

To determine the concentrations of *NF*-*κB* associated with the values of *k*_3_ and *k*_12_ that define the two lines of the cusp bifurcation diagram (Figure 13a), we constructed a 3D plot with an additional axis representing *NF*-*κB* (Figure 14a). Along each line of the cusp bifurcation diagram, the combined variation of *k*_3_ and *k*_12_ keeps the levels of *NF*-*κB* associated with each line unchanged. This behavior arises from the role of *p*65 in the reactions associated with *k*_3_ and *k*_12_. Although an increase in *k*_3_ reduces *p*65 levels through the reaction *p*65 → 0, and consequently decreases the availability of reactants in the reaction *p*50 + *p*65 → *NFκB*, this effect is compensated by the simultaneous increase in *k*_12_. These compensatory effects result in preserved levels of *NF*-*κB*.

**Fig. 14:**
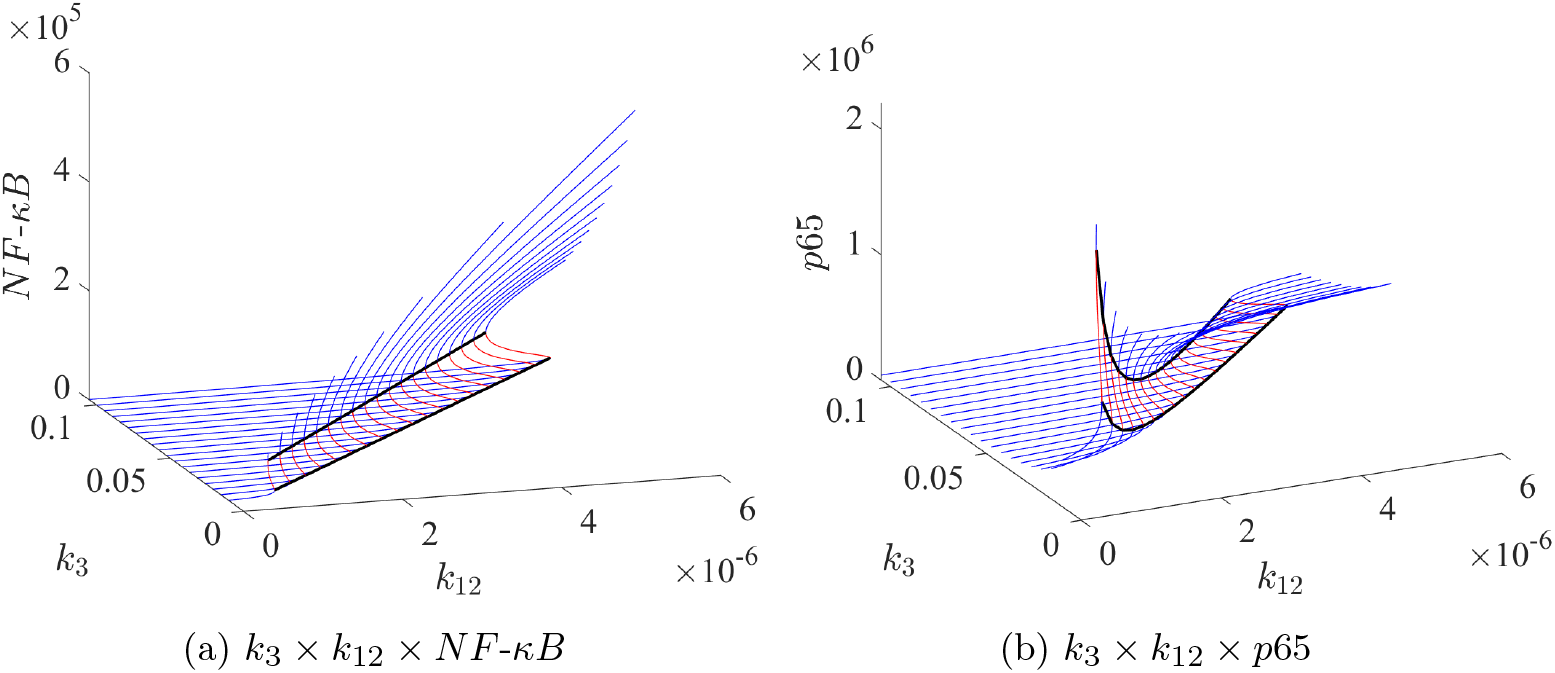
Three-dimensional bifurcation diagrams with free parameters *k*_3_ and *k*_12_. Subfigure (A) shows the behavior of *NF*-*κB*, while subfigure (B) depicts the behavior of *p*65.

Interestingly, for *p*65 levels, the combined variation of *k*_3_ and *k*_12_ along both lines of the cusp diagram produces a reinforcing rather than compensatory effect. Since both reactions associated with *k*_3_ and *k*_12_ reduce *p*65 levels, changing both parameters simultaneously results in a strong nonlinear reduction in *p*65 concentrations (Figure 14b).

Building on the observations regarding *NF*-*κB* and *p*65, the behavior of the remaining variables along the bifurcation lines can be understood from the stationary conditions of the GRN (1). Specifically, the concentrations of *N*0*p*50, *RNAp*50, *N*0*p*65, and *RNAp*65 remain constant, as each of these variables depends either on *NF*-*κB* or on other invariants along the lines (Figure S17). For example, the stationary condition for *N*0*p*50 is

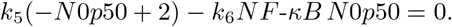

Since *k*_5_ and *k*_6_ are constants and *NF*-*κB* does not vary along the lines, *N*0*p*50 remains invariant. Similarly, for *RNAp*50 we have

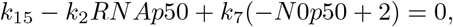

which implies that *RNAp*50 is also constant, as *N*0*p*50 does not change. The same reasoning applies to *N*0*p*65 and *RNAp*65, which are directly linked to these invariants.

For *p*50, the stationary condition reads

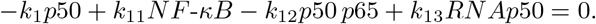

Here, *p*50 depends explicitly on both *NF*-*κB* and *p*65, as well as *RNAp*50. Along the bifurcation lines, *NF*-*κB* and *RNAp*50 remain constant. Increasing *k*_3_ reduces the levels of *p*65, through the association reaction *p*50 + *p*65 → *NF*-*κB*, this reduction in *p*65 would tend to decrease *NFκB* and, consequently, lower *p*50. However, a concurrent increase in *k*_12_ compensates for the drop in *p*65 by sustaining *NF*-*κB* levels. Under these conditions, the stationary balance above ensures that *p*50 remains effectively invariant despite the reduction in *p*65.

Since the levels of the *NF*-*κB* subunits, *p*65 and *p*50, strongly depends on its corresponding RNAs, we next turn our attention to the parameters *k*_7_ and *k*_10_, which correspond to the synthesis of *RNAp*50 (*N*1*p*50 → *N*1*p*50 + *RNAp*50) and *RNAp*65 (*N*1*p*65 → *N*1*p*65 + *RNAp*65), respectively. In this case, the bifurcation diagram exhibits two cusp points, (8.0877, 168.6359) and (86.8949, 16.8721), connected by two saddle-node bifurcation curves (Figure 15a). The bistable region delimited by these curves is a bounded set. Outside this region the system exhibits monostable behavior. Differently from the previous case, there are some intervals (*k*_7_ *<* 16.8721, *k*_7_ *>* 86.8949, *k*_10_ *<* 8.0877 and *k*_10_ *>* 168.6359) where a frontier between the HER2+ and TNBC regions is not well defined. To better characterize the subtypes associated to these regions, we built a 3D plot including the *NF*-*κB* axis, Figure 15b, which shows that within the closed region, the bifurcation diagram exhibits an S-shaped curve, reflecting the bistable behavior.

**Fig. 15:**
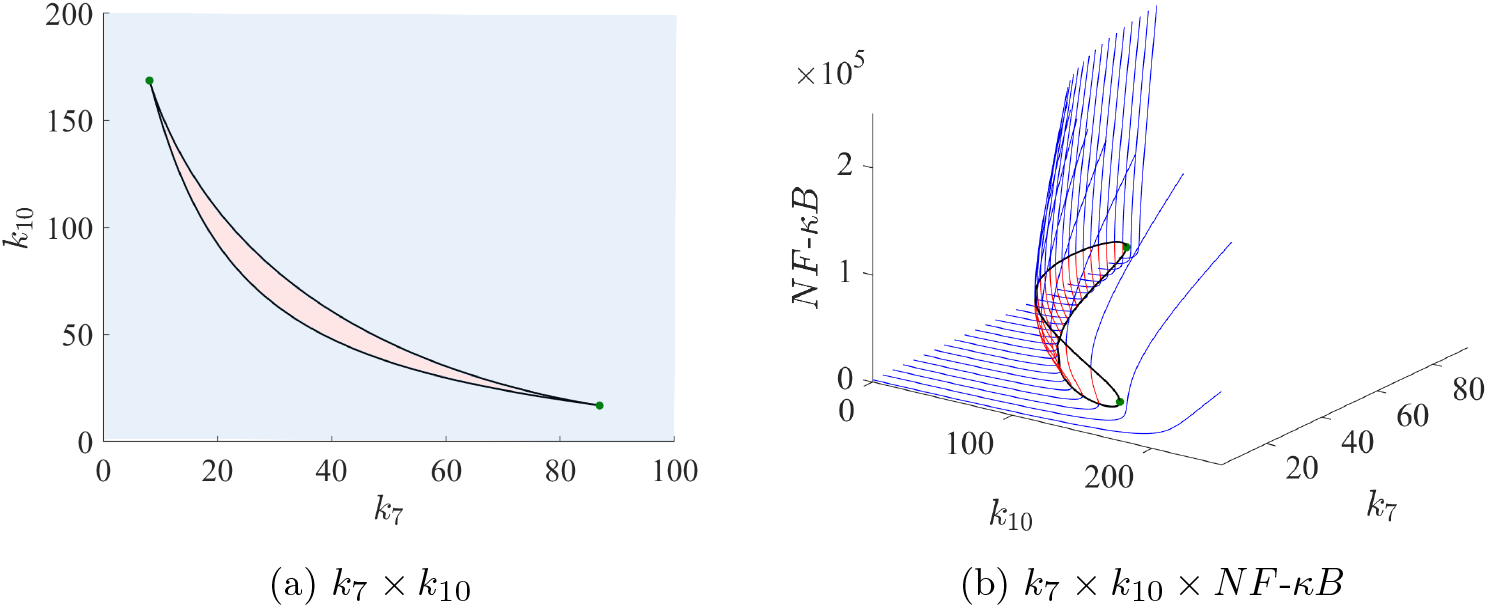
Bifurcation diagrams with free parameters *k*_7_ and *k*_10_. Subfigure (A) shows the bifurcation diagram in the parameter space: the green dots correspond to cusp points, the red region represents bistability, and the blue region represents monostability. Subfigure (B) presents the three-dimensional bifurcation diagram including the variable *NF*-*κB*.

To better visualize the differences between each region in Figure 15b, we plot three representative *NF*-*κB* level curves for different values of *k*_7_ in Figure 16. For *k*_7_ *<* 8.0877, as shown in Figure 16a, *NF*-*κB* increases continuously with *k*_10_. For the chosen value *k*_7_ = 3.706962815, when *k*_10_ ≲ 200, *NF*-*κB* levels are compatible with the HER2+ subtype, whereas for higher *k*_10_ values, they progressively increase toward TNBC-compatible levels.

**Fig. 16:**
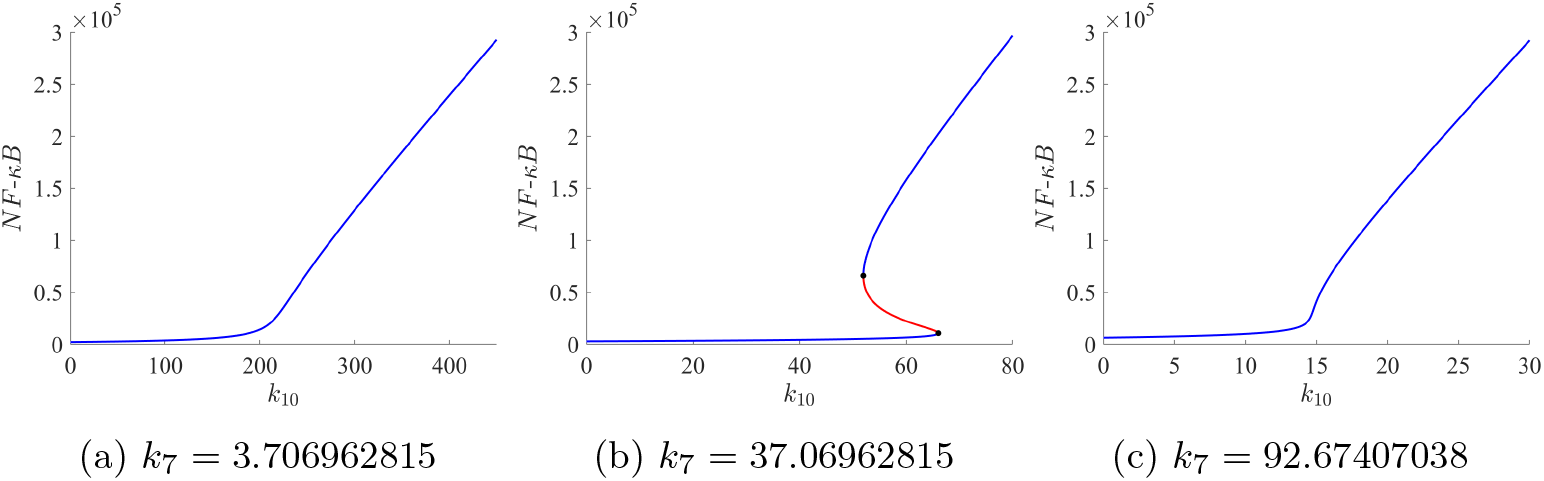
Bifurcation diagrams with only *k*_10_ free for different values of *k*_7_.

For *k*_7_ within the interval (8.0877, 86.8949) the bifurcation diagram with respect to *k*_10_ (Figure 16b) exhibits an S-shaped curve, indicating bistability. For *k*_7_ *>* 86.8949, *NF*-*κB* increases continuously with *k*_10_. In the specific case *k*_7_ = 92.67407038, *NF*-*κB* levels correspond to the HER2+ subtype when *k*_10_ ≲ 15, shifting toward TNBC-like levels as *k*_10_ increases (Figure 16c).

It is worth highlighting that the HER2+ to TNBC transition within the bistable region occurs under fixed parameter values, as the result of stochastic variations in the proteins and RNAs levels (see Lopes et al. (2025)). Consequently, outside this region, where only one stationary point exists, a HER2+ cell cannot expontaneously progress to the TNBC subtype, since no significative variation in parameter values are expected. However, under special conditions like mutations, the HER2+ to TNBC transition could happen, even outside the bistable regions.

This study reveals that the geometry of the bistable region is strongly parameter-dependent. For *k*_3_ and *k*_12_, the region is unbounded, with a single cusp point, whereas for *k*_7_ and *k*_10_ it is limited, with two cusp points. These results highlight how different parameter combinations can generate qualitatively distinct behaviors, emphasizing the system’s sensitivity to parametric changes and the importance of understanding these regions to interpret potential transitions between biological states, such as HER2+ and TNBC.

## 3 Materials and methods

### 3.1 Saddle-node bifurcation

To rigorously understand the qualitative changes observed in the system dynamics as parameters vary, we first recall the mathematical conditions under which a saddle-node bifurcation occurs. Consider a smooth one-parameter family of vector fields:

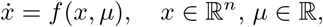

and suppose that for some (*x*_0_, *µ*_0_), we have *f* (*x*_0_, *µ*_0_) = 0, that is, *x*_0_ is an stationary point at the parameter value *µ*_0_. Let *A* = *D*_*x*_*f* (*x*_0_, *µ*_0_) be the Jacobian matrix evaluated at this point.

We say that the system undergoes a saddle-node bifurcation at (*x*_0_, *µ*_0_) if the following conditions are satisfied:

1. The matrix *A* has a simple eigenvalue equal to zero, and all other eigenvalues have nonzero real part;
2. Let *v* be the right eigenvector (*Av* = 0) and *w* the left eigenvector (*A*^⊤^*w* = 0) associated with the zero eigenvalue. Then, the nondegeneracy and transversality conditions are, respectively,

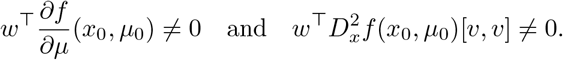

These conditions ensure the local existence of a smooth curve of equilibria tangent to the parameter axis at (*x*_0_, *µ*_0_). As the parameter *µ* crosses the bifurcation value *µ*_0_, the system transitions from having no stationary point near *x*_0_ to having two stationary points: one stable and one unstable (or vice versa). This behavior characterizes a saddle-node bifurcation.

Figure 17 depicts the characteristic behavior of a saddle-node bifurcation in the one-dimensional system 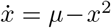, with (*x*_0_, *µ*_0_) = (0, 0) representing the saddle-node point.

**Fig. 17:**
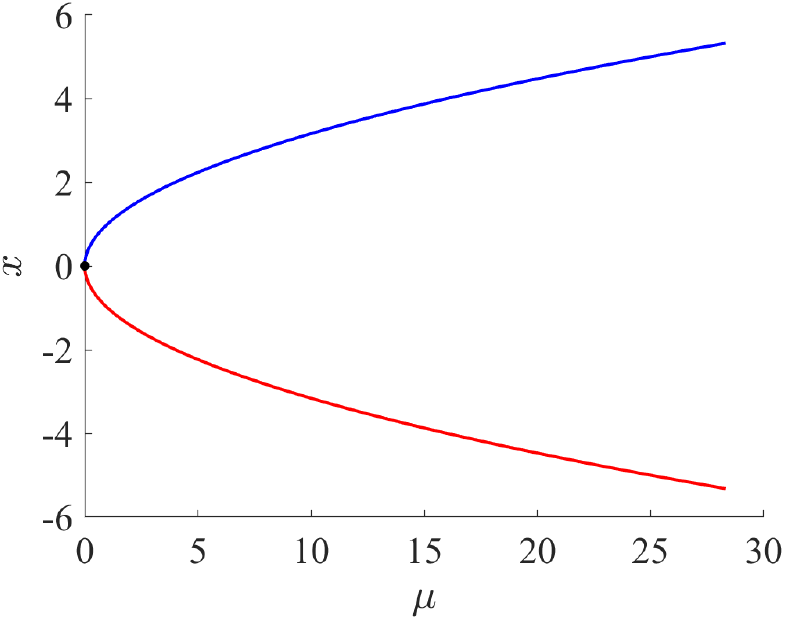
Saddle-node bifurcation diagram. The blue curve represents stable points, the red curve represents unstable points, and the black point represents the saddle-node point.

### 3.2 Cusp bifurcation

To characterize the qualitative changes in the system as two parameters vary, it is useful to recall the mathematical conditions for a cusp bifurcation. Consider a smooth two-parameter family of vector fields:

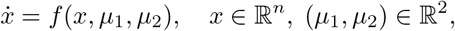

and suppose that at some point 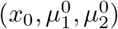, we have 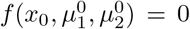, that is, *x*_0_ is stationary point at the parameter values 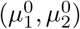. Let 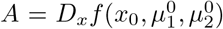 be the Jacobian matrix evaluated at this point.

The system undergoes a cusp bifurcation at 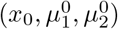 if the following conditions hold:

1. The Jacobian matrix *A* has a simple eigenvalue equal to zero, and all other eigenvalues have nonzero real part;
2. Let *v* be the right eigenvector and *w* the left eigenvector associated with the zero eigenvalue. Then the following nondegeneracy conditions are satisfied:

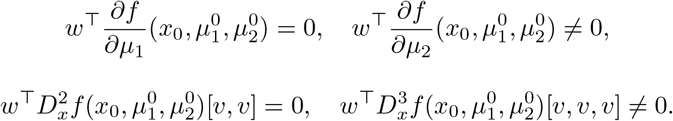

Specifically, a cusp bifurcation arises when two saddle-node bifurcation points merge and annihilate each other as the parameters vary. In parameter space, this gives rise to a bifurcation curve forming a cusp, which separates regions with different numbers of equilibria.

### 3.3 MatCont

The bifurcation analyses were performed using MatCont (version 7.6) A. Dhooge and W. Govaerts and Yu.A. Kuznetsov and H.G.E. Meijer and B. Sautois (2003– 2023), a *MATLAB* -based package for numerical continuation and stability analysis of dynamical systems defined by ordinary differential equations. The mathematical model considered is described in Section 2, and the numerical continuation was carried out with respect to different parameters of the model.

In the `Continuer` tab of MatCont, the following settings were adopted for the bifurcation analyses:

1. `InitStepsize` = 100
2. `MinStepsize` = 100
3. `MaxStepsize` = 1000
4. `MaxNewtonIters` = 3
5. `MaxCorrIters` = 10
6. `MaxTestIters` = 10
7. `VarTolerance` = 10^−6^
8. `FunTolerance` = 10^−6^
9. `TestTolerance` = 10^−5^
10. `Adapt` = 3
11. `MaxNumPoints` = varied depending on the case
12. `CheckClosed` = 50
13. `Jacobian Increment` = 10^−5^

All computations were carried out in *MATLAB* R2024a (Update 7) running on Windows 11, with MatCont version 7.6.

### 3.4 COPASI

Stochastic simulations were performed using *COPASI* (version 4.45) Hoops et al. (2006), a software tool for modeling and simulation of biochemical networks. The simulations employed the *τ* -Leap stochastic algorithm with the following settings: *ε* = 0.001, maximum number of internal steps = 1.215752192 × 10^7^, and a random seed fixed to 1.

Each simulation was run for an initial period of 262,800 minutes (approximately 6 months), with outputs recorded every 1 minute. If no transition occurred within this period, the simulation was extended iteratively in additional 6-month intervals until the transition was observed. If the transition did not occur within 20 years of simulated time, the run was terminated. Initial conditions were set at the steady state of the deterministic model whenever required.

## 4 Discussion

A central question in the study of transitions between stable states in nonlinear dynamical systems is which dynamical features effectively control transition probabilities. In many theoretical frameworks, this control is often interpreted in terms of potential depth. However, in non-conservative systems, such as gene regulatory networks, potential functions are generally non-unique and do not uniquely determine the system?s dynamics, which limits their explanatory and predictive power.

Our results provide an alternative and more general perspective by showing that the phase-space distance between stationary states is a critical determinant of transition probabilities. Unlike potential-based measures, state-space distances are well defined in non-equilibrium systems and can, in principle, be estimated experimentally. In this context, the bifurcation structure of the system plays a key role in shaping how such distances emerge and constrain accessible transitions between stable states.

Importantly, the mere coexistence of multiple stable states is only a necessary, but not sufficient, condition for transitions to occur. The presence of two basins of attraction does not, by itself, imply that transitions between them are likely or even dynamically accessible on relevant timescales. Instead, transition probabilities are shaped by how these basins are arranged in phase space and, in particular, by the distance separating stable states from the intervening unstable states. From this perspective, phase-space distance acts as an effective control parameter, simultaneously constraining the likelihood, timescale, and variability of transitions. This form of control is inherently geometric rather than energetic, and does not rely on the existence of a well-defined potential function.

In addition to the role of phase-space distance, our previous analysis of the NF-*κ*B regulatory system Lopes et al. (2025) demonstrated that the local dynamical structure of the intervening unstable stationary state further constrains transition dynamics. Linear stability analysis revealed that this state possesses a single unstable eigendirection associated with a small positive eigenvalue, while all remaining directions are strongly contracting. This configuration generates a narrow one-dimensional escape channel embedded within a predominantly attracting local flow. Stochastic trajectories approaching this region are therefore funneled toward the unstable manifold, and once the vicinity of the saddle is reached, crossing occurs rapidly along this weakly unstable direction. In this sense, the unstable state functions as a geometric gateway between basins: the dominant constraint lies in reaching its neighborhood, whereas escape beyond it is dynamically swift. This mechanism reinforces the geometric character of transition control in non-conservative systems, where the local eigenspectrum and the global phase-space distances jointly determine accessibility, irreversibility, and variability of phenotypic transitions.

Within this geometric framework, our bifurcation analysis reveals a marked asymmetry in the sensitivity of the two stable states to parameter variations. Across all model parameters, the HER2+ branch consistently exhibits low sensitivity, with NF-*κ*B levels remaining relatively invariant even under substantial parameter changes. This behavior suggests that, in this regime, NF-*κ*B effectively buffers intrinsic parameter variability in upstream processes, acting as a dynamical filter that suppresses the transmission of such variability across the network.

In contrast, the TNBC branch displays pronounced sensitivity to the same parameter variations, with small parameter changes leading to large variations in NF-*κ*B levels. In this regime, intrinsic parameter variability is not attenuated but instead amplified by the network architecture, resulting in heightened variability of the system?s state. This intrinsic dynamical instability provides a natural interpretation for the extensive intra- and inter-tumoral variability observed in TNBC, as well as for the large number of molecular subclassifications proposed for this subtype Lehmann et al. (2016); Burstein et al. (2015). From this perspective, the clinical variability associated with TNBC emerges as a direct consequence of its underlying dynamical organization, rather than solely from genetic or environmental differences.

Beyond sensitivity at fixed parameter values, our two-parameter bifurcation analysis shows that the global geometry of the bistable region itself depends critically on the choice of regulatory mechanisms being perturbed. In particular, different parameter combinations give rise to qualitatively distinct cusp geometries, resulting in either unbounded or limited bistable regions in parameter space. Unbounded regions imply that bistability can persist over wide ranges of parameter variation, allowing stable states to remain dynamically accessible despite substantial intrinsic variation. In contrast, bounded bistable regions impose strict limits on the parameter ranges compatible with multistability, beyond which one of the stable states disappears altogether.

From a dynamical perspective, these geometric differences introduce an additional level of constraint on phenotypic transitions. While phase-space distance governs transition probabilities within a bistable regime, the cusp geometry determines whether such a regime exists at all under parameter variation. As a result, the accessibility of transitions is controlled not only by distances between states, but also by the global structure of the bifurcation set that organizes the emergence and destruction of attractor basins. This highlights the role of cusp bifurcations as higher-order organizing centers that delimit the regions of phenotypic plasticity in non-conservative regulatory networks.

Taken together, our results establish a geometric framework in which transitions in non-conservative regulatory networks are controlled by a hierarchy of structural features, including phase-space distances between stable states, their sensitivity to intrinsic parameter perturbations, and the global organization imposed by cusp bifurcations. Importantly, this framework does not rely on system-specific molecular details and is therefore expected to apply broadly to gene regulatory networks and other nonlinear dynamical systems operating far from equilibrium.

The present analysis focuses on how distances and bifurcation geometry constrain transition probabilities and times under parameter variation. An additional dynamical aspect that remains outside the scope of this work concerns the observation that identical phase-space distances between stationary states do not necessarily produce identical transition-time distributions. As evidenced by the transition-time statistics reported here, different kinetic perturbations that yield the same separation between states can still lead to distinct stochastic escape dynamics. This indicates that, beyond distance, further structural properties of the underlying flow influence transition statistics. Characterizing these additional geometric?dynamical factors represents a natural extension of the present framework and will be addressed in subsequent work.

## Supplementary information

Supplementary material accompanies this article and contains additional figures and tables supporting the results.

## Data Availability

No external datasets were used in this study. The data supporting the findings were generated through computational simulations and are available from the corresponding author upon reasonable request.

## Acknowledgements

The authors acknowledge the Laboratório Nacional de Computação Científica (LNCC/MCTI, Brazil) for providing access to the Santos Dumont supercomputer and a MATLAB license. The authors also thank the High Performance Computer Center (NACAD) - COPPE/Federal University of Rio de Janeiro (UFRJ), which has supported this research. Website: http://nacad.ufrj.br. The authors thank Alfonso Landeros for carefully reading the manuscript and for valuable suggestions that helped improve the paper.

## References

A. Dhooge and W. Govaerts and Yu.A. Kuznetsov and H.G.E. Meijer and B. Sautois: Matcont: A MATLAB package for numerical bifurcation analysis of ODEs. Disponível em: http://www.matcont.ugent.be/ (2003–2023)

Burstein, M.D., Tsimelzon, A., Poage, G.M., Covington, K.R., Contreras, A., Fuqua, S.A., Savage, M.I., Osborne, C.K., Hilsenbeck, S.G., Chang, J.C., et al.: Comprehensive genomic analysis identifies novel subtypes and targets of triple-negative breast cancer. Clinical Cancer Research 21(7), 1688–1698 (2015)

Chung, W., Eum, H.H., Lee, H.-O., Lee, K.-M., Lee, H.-B., Kim, K.-T., Ryu, H.S., Kim, S., Lee, J.E., Park, Y.H., et al.: Single-cell rna-seq enables comprehensive tumour and immune cell profiling in primary breast cancer. Nature communications 8(1), 15081 (2017)

Espenson, J.H.: Chemical Kinetics and Reaction Mechanisms. McGraw-Hill, ??? (1995)

Huang, S., Ernberg, I., Kauffman, S.: Cancer attractors: a systems view of tumors from a gene network dynamics and developmental perspective. Seminars in Cell & Developmental Biology 20(7), 869–876 (2009)

Hsu, C., Jaquet, V., Gencoglu, M., Becskei, A.: Protein dimerization generates bistability in positive feedback loops. Cell reports 16(5), 1204–1210 (2016)

Hoops, S., Sahle, S., Gauges, R., Lee, C., Pahle, J., Simus, N., Singhal, M., Xu, L., Mendes, P., Kummer, U.: Copasi: a complex pathway simulator. Bioinformatics 22(24), 3067–3074 (2006) 10.1093/bioinformatics/btl485

Kauffman, S.: Differentiation of malignant to benign cells. Journal of theoretical biology 31(3), 429–451 (1971)

Lehmann, B.D., Jovanović, B., Chen, X., Estrada, M.V., Johnson, K.N., Shyr, Y., Moses, H.L., Sanders, M.E., Pietenpol, J.A.: Refinement of triple-negative breast cancer molecular subtypes: implications for neoadjuvant chemotherapy selection. PloS one 11(6), 0157368 (2016)

Lopes, F., Pires, B.R., Lima, A.A., Binato, R., Abdelhay, E.: Nf-κb epigenetic attractor landscape drives breast cancer heterogeneity. npj Systems Biology and Applications 11(1), 135 (2025)

Lüönd, F., Tiede, S., Christofori, G.: Breast cancer as an example of tumour heterogeneity and tumour cell plasticity during malignant progression. British journal of cancer 125(2), 164–175 (2021)

Lopes, F.J., Vieira, F.M., Holloway, D.M., Bisch, P.M., Spirov, A.V.: Spatial bistability generates hunchback expression sharpness in the drosophila embryo. PLoS computational biology 4(9), 1000184 (2008)

Maisonneuve, P., Disalvatore, D., Rotmensz, N., Curigliano, G., Colleoni, M., Dellapasqua, S., Pruneri, G., Mastropasqua, M.G., Luini, A., Bassi, F., et al.: Proposed new clinicopathological surrogate definitions of luminal a and luminal b (her2-negative) intrinsic breast cancer subtypes. Breast Cancer Research 16(3), 65 (2014)

Pahl, H.L.: Activators and target genes of rel/nf-κb transcription factors. Oncogene 18(49), 6853–6866 (1999)

Pavitra, E., Kancharla, J., Gupta, V.K., Prasad, K., Sung, J.Y., Kim, J., Tej, M.B., Choi, R., Lee, J.-H., Han, Y.-K., et al.: The role of nf-κb in breast cancer initiation, growth, metastasis, and resistance to chemotherapy. Biomedicine & pharmacotherapy 163, 114822 (2023)

Pires, B.R., Mencalha, A.L., Ferreira, G.M., De Souza, W.F., Morgado-Díaz, J.A., Maia, A.M., Corrêa, S., Abdelhay, E.S.: Nf-kappab is involved in the regulation of emt genes in breast cancer cells. PloS one 12(1), 0169622 (2017)

Sung, H., Ferlay, J., Siegel, R.L., Laversanne, M., Soerjomataram, I., Jemal, A., Bray, F.: Global cancer statistics 2020: Globocan estimates of incidence and mortality worldwide for 36 cancers in 185 countries. CA: a cancer journal for clinicians 71(3), 209–249 (2021)

Sung, M.-H., Li, N., Lao, Q., Gottschalk, R.A., Hager, G.L., Fraser, I.D.: Switching of the relative dominance between feedback mechanisms in lipopolysaccharide-induced nf-κb signaling. Science signaling 7(308), 6–6 (2014)

Waddington, C.: The Strategy of the Genes. Allen. Unwin, London (1957)

Wu, S.Z., Al-Eryani, G., Roden, D.L., Junankar, S., Harvey, K., Andersson, A., Thennavan, A., Wang, C., Torpy, J.R., Bartonicek, N., et al.: A single-cell and spatially resolved atlas of human breast cancers. Nature genetics 53(9), 1334–1347 (2021)

Wang, W., Nag, S.A., Zhang, R.: Targeting the nfκb signaling pathways for breast cancer prevention and therapy. Current medicinal chemistry 22(2), 264–289 (2015)

