## supplementary material for "Phase-space distance between stationary states modulates phenotypic plasticity in breast cancer"

[1,2,3] Developmental Biology and Dynamical Systems Group, Universidade Federal do Rio de Janeiro, Campus Duque de Caxias Professor Geraldo Cidade; Duque de Caxias, Brazil.

### Reactions

| **Reaction** | **Kinetic constant** | Unit |
|---|---|---|
| $p50 \to 0$ | $k_1$ | $\mathrm{min}^{-1}$ |
| $RNAp50 \to 0$ | $k_2$ | $\mathrm{min}^{-1}$ |
| $p65 \to 0$ | $k_3$ | $\mathrm{min}^{-1}$ |
| $RNAp65 \to 0$ | $k_4$ | $\mathrm{min}^{-1}$ |
| $N1p50 \to N0p50 + NF\text{-}\kappa B$ | $k_5$ | $\mathrm{min}^{-1}$ |
| $N0p50 + NF\text{-}\kappa B \to N1p50$ | $k_6$ | $(\# \cdot \mathrm{min})^{-1}$ |
| $N1p50 \to N1p50 + RNAp50$ | $k_7$ | $\mathrm{min}^{-1}$ |
| $N1p65 \to N0p65 + NF\text{-}\kappa B$ | $k_8$ | $\mathrm{min}^{-1}$ |
| $N0p65 + NF\text{-}\kappa B \to N1p65$ | $k_9$ | $(\# \cdot \mathrm{min})^{-1}$ |
| $N1p65 \to N1p65 + RNAp65$ | $k_{10}$ | $\mathrm{min}^{-1}$ |
| $NF\text{-}\kappa B \to p50 + p65$ | $k_{11}$ | $\mathrm{min}^{-1}$ |
| $p50 + p65 \to NF\text{-}\kappa B$ | $k_{12}$ | $(\# \cdot \mathrm{min})^{-1}$ |
| $RNAp50 \to RNAp50 + p50$ | $k_{13}$ | $\mathrm{min}^{-1}$ |
| $RNAp65 \to RNAp65 + p65$ | $k_{14}$ | $\mathrm{min}^{-1}$ |
| $0 \to RNAp50$ | $k_{15}$ | $\# \cdot \mathrm{min}^{-1}$ |
| $0 \to RNAp65$ | $k_{16}$ | $\# \cdot \mathrm{min}^{-1}$ |

Table S1: List of biochemical reactions and their corresponding kinetic rate constants, including units, defining the gene regulatory network shown in Figure 1. The symbol # denotes molecule copy number.

### Bifurcation diagrams with a free parameter

| $k$'s | Reaction | Bistability | HER2+ | TNBC |
|---|---|---|---|---|
| $k_1$ | $p50 \to 0$ | $(0.028, 0.032)$ | $(0.032, 0.043)$ | $(0.024, 0.028)$ |
| $k_2$ | $RNAp50 \to 0$ | $(0.093, 0.109)$ | $(0.109, 0.145)$ | $(0.082, 0.093)$ |
| $k_3$ | $p65 \to 0$ | $(0.048, 0.056)$ | $(0.056, 0.0655)$ | $(0.0403, 0.048)$ |
| $k_4$ | $RNAp65 \to 0$ | $(0.099, 0.115)$ | $(0.115, 0.148)$ | $(0.0802, 0.099)$ |
| $k_5$ | $N1p50 \to N0p50 + NF\text{-}\kappa B$ | $(70230.135, 94576.068)$ | $(94576.068, 114480)$ | $(45192, 70230.135)$ |
| $k_6$ | $N0p50 + NF\text{-}\kappa B \to N1p50$ | $(0.791, 1.066)$ | $(0.6316, 0.791)$ | $(1.066, 1.1782)$ |
| $k_7$ | $N1p50 \to N1p50 + RNAp50$ | $(34.321, 42.928)$ | $(26.2791, 34.321)$ | $(42.928, 46.1260)$ |
| $k_8$ | $N1p65 \to N0p65 + NF\text{-}\kappa B$ | $(87.845, 118.548)$ | $(118.548, 127.2987)$ | $(72.6103, 87.845)$ |
| $k_9$ | $N0p65 + NF\text{-}\kappa B \to N1p65$ | $(0.00079, 0.0011)$ | $(0.00066, 0.00079)$ | $(0.0011, 0.0012)$ |
| $k_{10}$ | $N1p65 \to N1p65 + RNAp65$ | $(51.956, 66.055)$ | $(41.7493, 51.956)$ | $(66.055, 75.1749)$ |
| $k_{11}$ | $NF\text{-}\kappa B \to p50 + p65$ | $(1.556, 1.809)$ | $(1.809, 2.0817)$ | $(1.3884, 1.556)$ |
| $k_{12}$ | $p50 + p65 \to NF\text{-}\kappa B$ | $(2.498{\times}10^{-6}, 2.905{\times}10^{-6})$ | $(2.348{\times}10^{-6}, 2.498{\times}10^{-6})$ | $(2.905{\times}10^{-6}, 3.12{\times}10^{-6})$ |
| $k_{13}$ | $RNAp50 \to RNAp50 + p50$ | $(8.798, 10.23)$ | $(7.9287, 8.798)$ | $(10.23, 11.1441)$ |
| $k_{14}$ | $RNAp65 \to RNAp65 + p65$ | $(42.162, 49.024)$ | $(40.1739, 42.162)$ | $(49.024, 51.7269)$ |
| $k_{15}$ | $0 \to RNAp50$ | $(3.732, 7.517)$ | $(1.7303, 3.732)$ | $(7.517, 9.7718)$ |
| $k_{16}$ | $0 \to RNAp65$ | $(5.987, 11.047)$ | $(2.6464, 5.987)$ | $(11.047, 13.0451)$ |

Table S2: Bistability and monostability intervals for each parameter. The first column represents the parameters, the second the corresponding reactions, the third represents the bistability interval for each parameter, and the fourth and fifth columns represent the monostability intervals, identifying which subtype corresponds to HER2+ or TNBC.

| Variables | Δ HER2+ | Δ TNBC | Δ Total | % HER2+ | % TNBC |
|---|---|---|---|---|---|
| $RNAp50$ | 65.70 | 216.15 | 507.10 | 12.96 | 42.63 |
| $RNAp65$ | 75.52 | 315.55 | 675.03 | 11.19 | 46.75 |
| $NF\text{-}\kappa B$ | 9602.79 | 169207.40 | 234536.93 | 4.09 | 72.15 |
| $N0p50$ | 0.18 | 0.60 | 1.39 | 12.96 | 42.63 |
| $N0p65$ | 0.14 | 0.61 | 1.29 | 11.19 | 46.75 |
| $p50$ | 31957.30 | 117365.70 | 208184.18 | 15.35 | 56.38 |
| $p65$ | 64652.90 | 270137.66 | 577890.91 | 11.19 | 46.75 |

Table S3: Variation of each species' copy number along the HER2+ and TNBC branches as a function of $k_1$ in the corresponding bifurcation diagrams. The columns are as follows: (1) variables; (2) absolute variation along the TNBC branch; (3) absolute variation along the HER2+ branch; (4) total absolute variation across the entire bifurcation diagram; (5-6) the percentage values, with column (4) representing 100%.

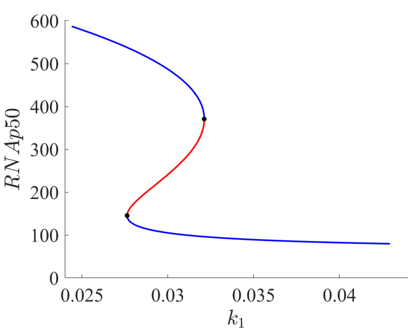


(a) $k_1 \times RNAp50$

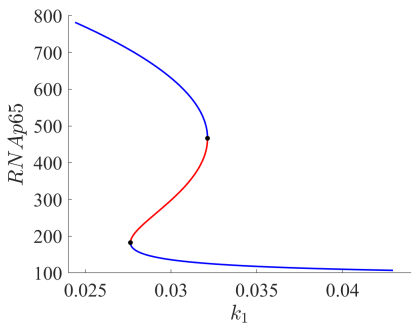


(b) $k_1 \times RNAp65$

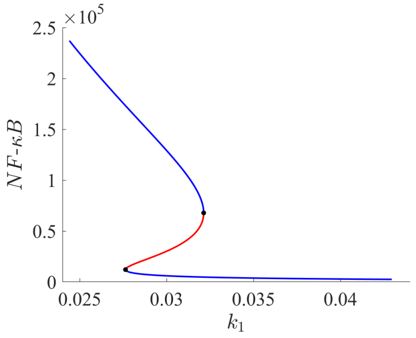


(c) $k_1 \times NF\text{-}\kappa B$

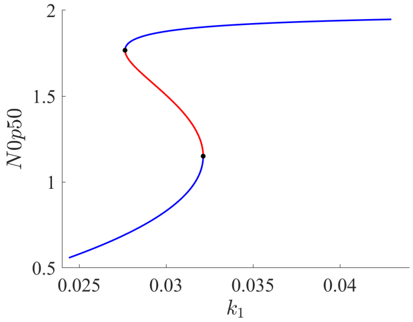


(d) $k_1 \times N0p50$

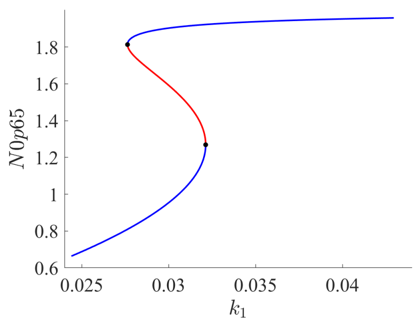


(e) $k_1 \times N0p65$

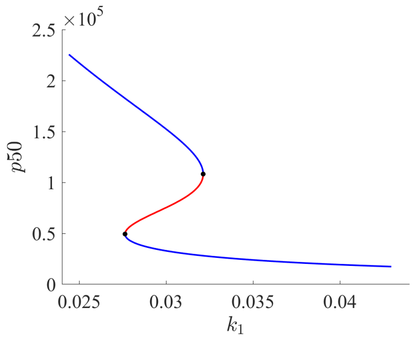


(f) $k_1 \times p50$

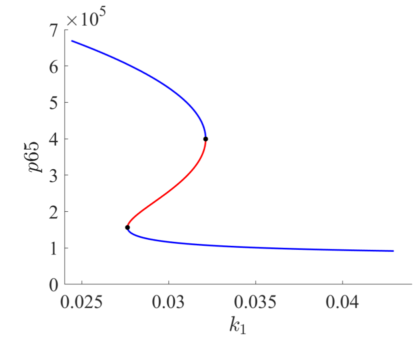


(g) $k_1 \times p65$

Figure S1: Bifurcation diagrams for all species with respect to the parameter $k_1$

| **Variables** | **TNBC** | **HER2+** | **Total** | **% TNBC** | **% HER2+** |
|---|---|---|---|---|---|
| $RNAp50$ | 377.03 | 102.66 | 668.78 | 56.38 | 15.35 |
| $RNAp65$ | 315.55 | 75.52 | 675.03 | 46.75 | 11.19 |
| $NF\text{-}\kappa B$ | 169207.21 | 9602.80 | 234536.75 | 72.15 | 4.09 |
| $N0p50$ | 0.59 | 0.18 | 1.39 | 42.63 | 12.96 |
| $N0p65$ | 0.61 | 0.14 | 1.29 | 46.75 | 11.19 |
| $p50$ | 117365.57 | 31957.37 | 208184.12 | 56.38 | 15.35 |
| $p65$ | 270137.50 | 64652.99 | 577890.84 | 46.75 | 11.19 |

Table S4: Variation of each species' copy number along the HER2+ and TNBC branches as a function of $k_2$ in the corresponding bifurcation diagrams. The columns are as follows: (1) variables; (2) absolute variation along the TNBC branch; (3) absolute variation along the HER2+ branch; (4) total absolute variation across the entire bifurcation diagram; (5-6) the percentage values, with column (4) representing 100%.

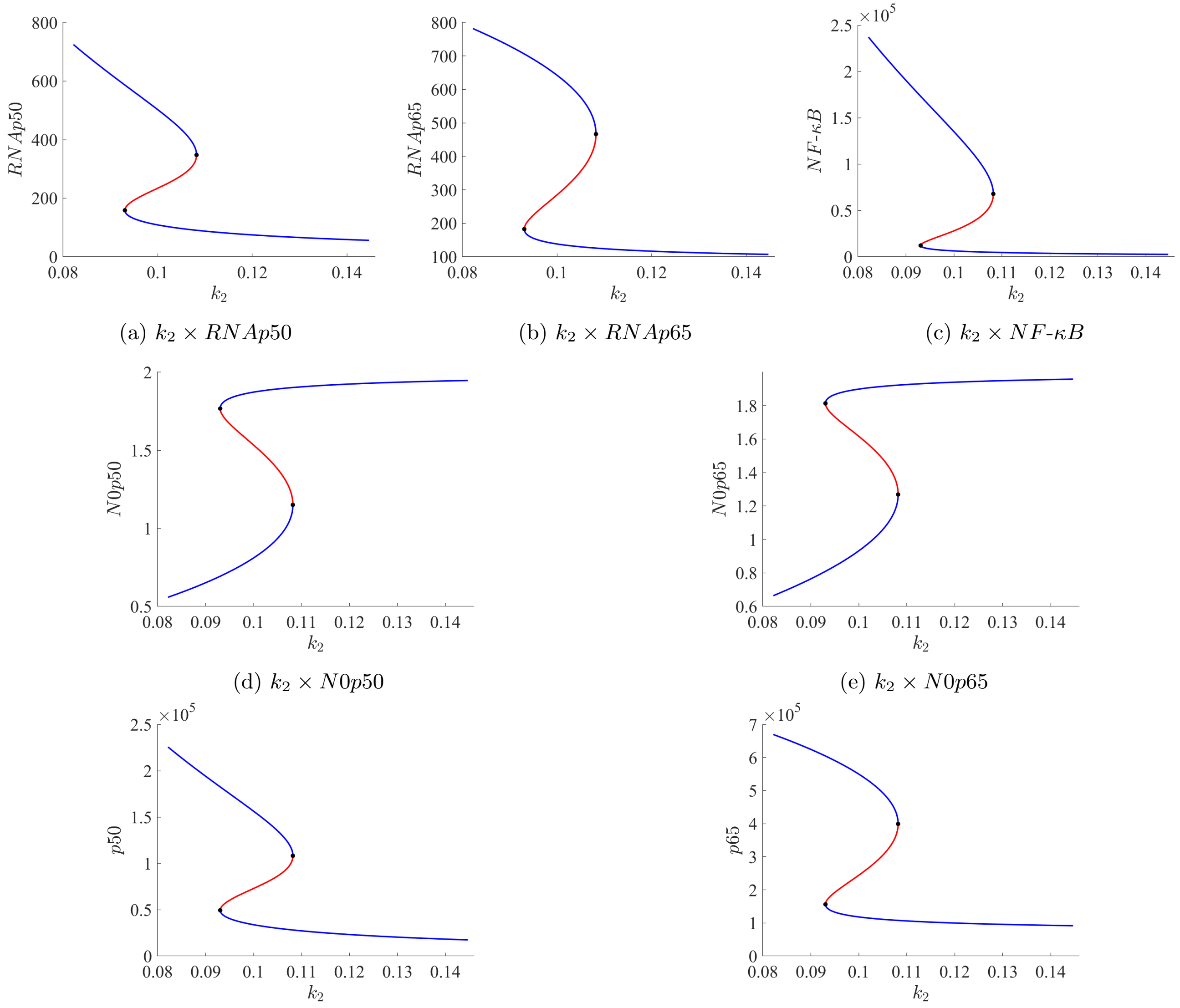


(a) $k_2 \times RNAp50$ (b) $k_2 \times RNAp65$ (c) $k_2 \times NF$-$\kappa B$

(d) $k_2 \times N0p50$ (e) $k_2 \times N0p65$

(f) $k_2 \times p50$ (g) $k_2 \times p65$

Figure S2: Bifurcation diagrams for all species with respect to the parameter $k_2$

| **Variables** | **TNBC** | **HER2+** | **Total** | **% TNBC** | **% HER2+** |
|---|---|---|---|---|---|
| $RNAp50$ | 240.97 | 64.36 | 530.58 | 45.42 | 12.13 |
| $RNAp65$ | 667.15 | 119.97 | 1025.61 | 65.05 | 11.70 |
| $NF$-$\kappa B$ | 214768.57 | 9424.62 | 279919.94 | 76.72 | 3.37 |
| $N0p50$ | 0.66 | 0.18 | 1.45 | 45.42 | 12.13 |
| $N0p65$ | 0.68 | 0.14 | 1.37 | 49.79 | 10.38 |
| $p50$ | 75012.15 | 20033.75 | 165165.27 | 45.42 | 12.13 |
| $p65$ | 571139.41 | 102708.90 | 878020.59 | 65.05 | 11.70 |

Table S5: Variation of each species' copy number along the HER2+ and TNBC branches as a function of $k_4$ in the corresponding bifurcation diagrams. The columns are as follows: (1) variables; (2) absolute variation along the TNBC branch; (3) absolute variation along the HER2+ branch; (4) total absolute variation across the entire bifurcation diagram; (5-6) the percentage values, with column (4) representing 100%.

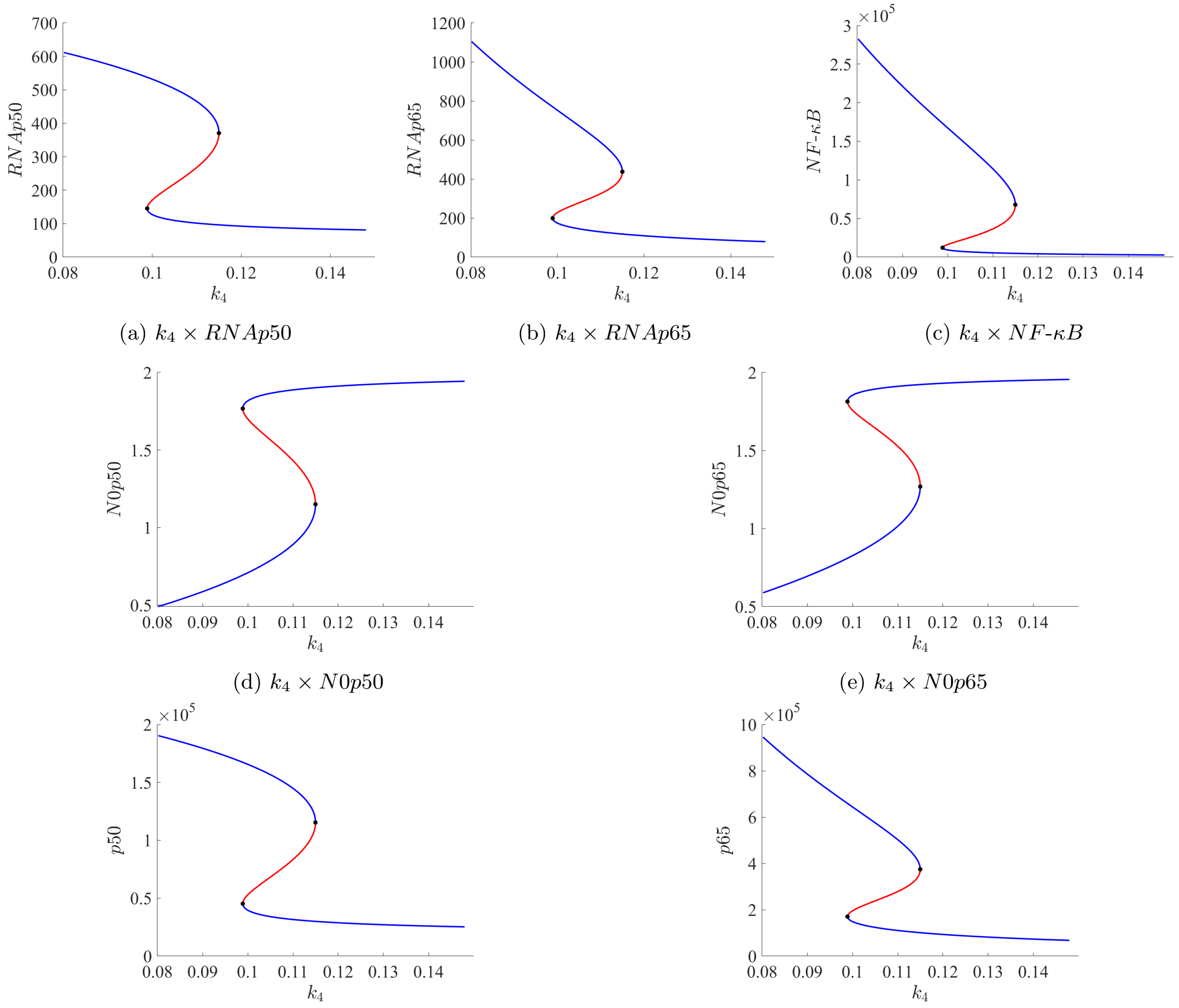


(a) $k_4 \times RNAp50$ (b) $k_4 \times RNAp65$ (c) $k_4 \times NF\text{-}\kappa B$

(d) $k_4 \times N0p50$ (e) $k_4 \times N0p65$

(f) $k_4 \times p50$ (g) $k_4 \times p65$

Figure S3: Bifurcation diagrams for all species with respect to the parameter $k_4$

| **Variables** | **TNBC** | **HER2+** | **Total** | **% TNBC** | **% HER2+** |
|---|---|---|---|---|---|
| $RNAp50$ | 284.65 | 68.48 | 560.18 | 50.81 | 12.23 |
| $RNAp65$ | 257.80 | 54.35 | 619.49 | 41.62 | 8.77 |
| $NF\text{-}\kappa B$ | 127088.00 | 6969.87 | 195592.10 | 64.98 | 3.56 |
| $N0p50$ | 0.78 | 0.19 | 1.53 | 50.81 | 12.23 |
| $N0p65$ | 0.49 | 0.10 | 1.19 | 41.62 | 8.77 |
| $p50$ | 88607.65 | 21318.80 | 174379.51 | 50.81 | 12.23 |
| $p65$ | 220701.90 | 46531.13 | 530340.57 | 41.62 | 8.77 |

Table S6: Variation of each species' copy number along the HER2+ and TNBC branches as a function of $k_5$ in the corresponding bifurcation diagrams. The columns are as follows: (1) variables; (2) absolute variation along the TNBC branch; (3) absolute variation along the HER2+ branch; (4) total absolute variation across the entire bifurcation diagram; (5-6) the percentage values, with column (4) representing 100%.

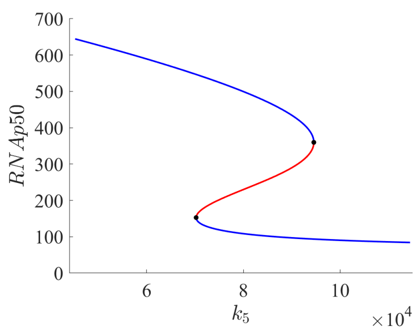


(a) $k_5 \times RNAp50$

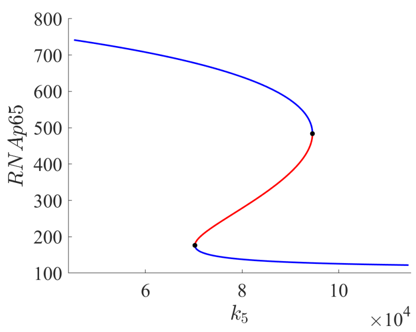


(b) $k_5 \times RNAp65$

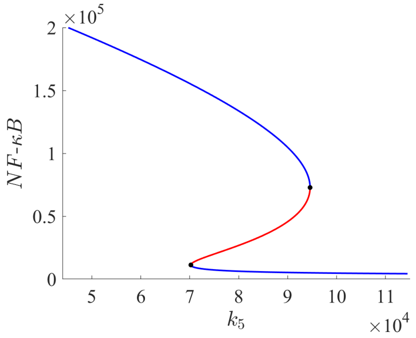


(c) $k_5 \times NF\text{-}\kappa B$

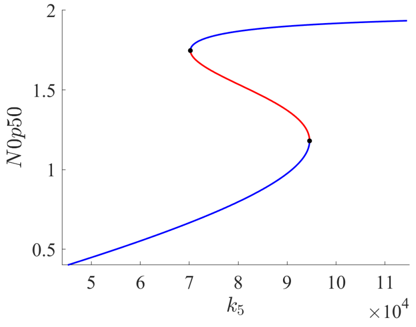


(d) $k_5 \times N0p50$

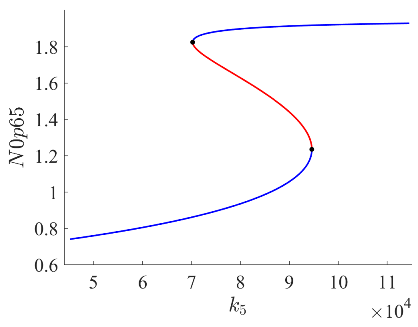


(e) $k_5 \times N0p65$

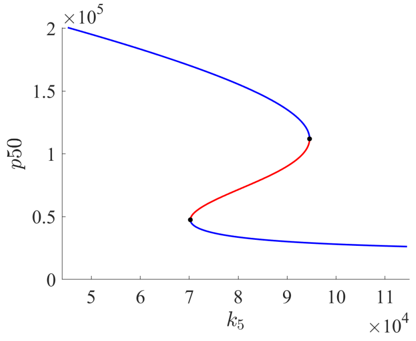


(f) $k_5 \times p50$

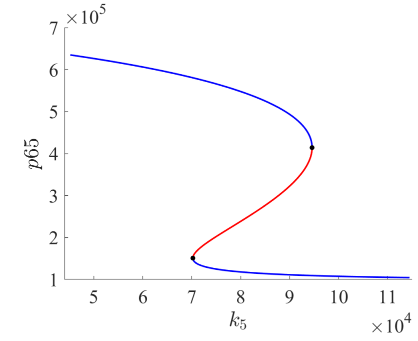


(g) $k_5 \times p65$

Figure S4: Bifurcation diagrams for all species with respect to the parameter $k_5$

| **Variables** | **TNBC** | **HER2+** | **Total** | **% TNBC** | **% HER2+** |
|---|---|---|---|---|---|
| $RNAp50$ | 215.29 | 69.75 | 492.08 | 43.75 | 14.17 |
| $RNAp65$ | 214.67 | 55.10 | 577.10 | 37.20 | 9.55 |
| $NF\text{-}\kappa B$ | 95199.77 | 7060.07 | 163794.06 | 58.12 | 4.31 |
| $N0p50$ | 0.59 | 0.19 | 1.35 | 43.75 | 14.17 |
| $N0p65$ | 0.41 | 0.11 | 1.11 | 37.20 | 9.55 |
| $p50$ | 67017.48 | 21711.16 | 153181.70 | 43.75 | 14.17 |
| $p65$ | 183774.79 | 47168.13 | 494050.46 | 37.20 | 9.55 |

Table S7: Variation of each species' copy number along the HER2+ and TNBC branches as a function of $k_6$ in the corresponding bifurcation diagrams. The columns are as follows: (1) variables; (2) absolute variation along the TNBC branch; (3) absolute variation along the HER2+ branch; (4) total absolute variation across the entire bifurcation diagram; (5-6) the percentage values, with column (4) representing 100%.

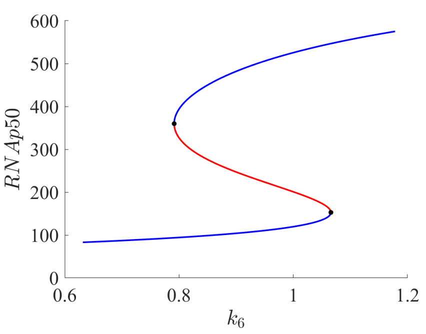


(a) $k_6 \times RNAp50$

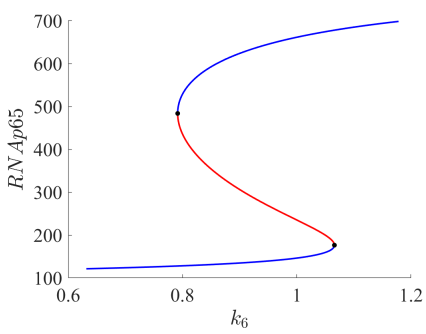


(b) $k_6 \times RNAp65$

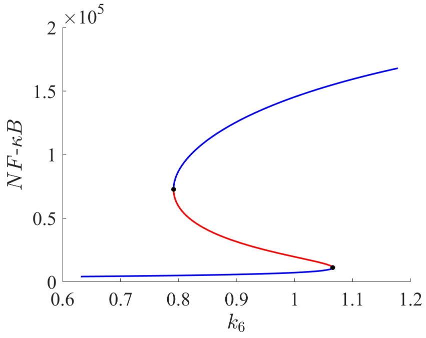


(c) $k_6 \times NF\text{-}\kappa B$

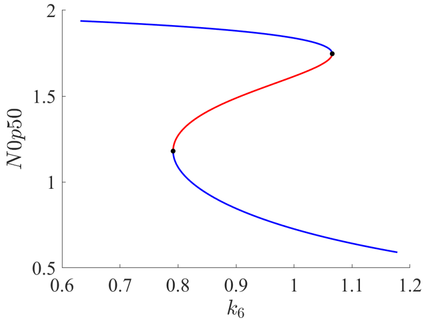


(d) $k_6 \times N0p50$

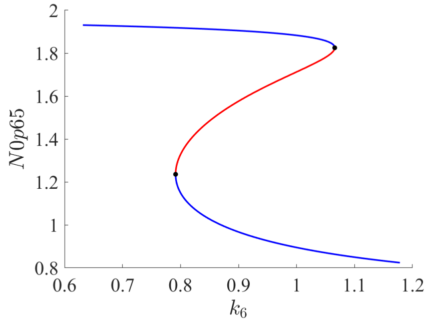


(e) $k_6 \times N0p65$

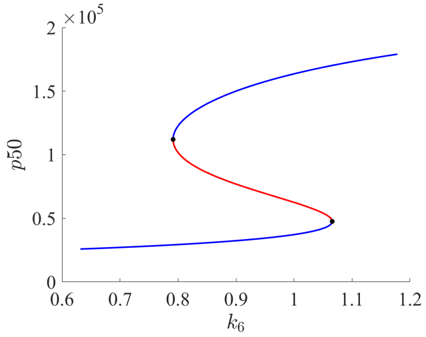


(f) $k_6 \times p50$

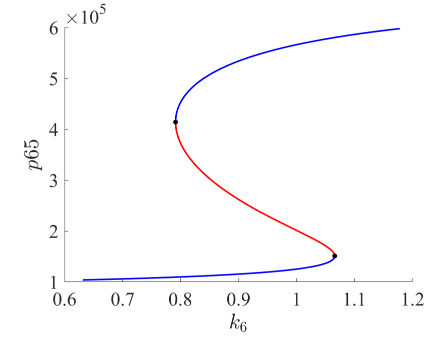


(g) $k_6 \times p65$

Figure S5: Bifurcation diagrams for all species with respect to the parameter $k_6$

| **Variables** | **TNBC** | **HER2+** | **Total** | **% TNBC** | **% HER2+** |
|---|---|---|---|---|---|
| $RNAp50$ | 366.08 | 66.94 | 627.04 | 58.38 | 10.68 |
| $RNAp65$ | 314.12 | 52.24 | 653.96 | 48.03 | 7.99 |
| $NF\text{-}\kappa B$ | 163904.54 | 6674.20 | 226043.61 | 72.51 | 2.95 |
| $N0p50$ | 0.5908 | 0.12 | 1.34 | 44.03 | 9.26 |
| $N0p65$ | 0.60 | 0.10 | 1.25 | 48.03 | 7.99 |
| $p50$ | 113957.07 | 20838.90 | 195191.51 | 58.38 | 10.68 |
| $p65$ | 268914.09 | 44723.95 | 559847.10 | 48.03 | 7.99 |

Table S8: Variation of each species' copy number along the HER2+ and TNBC branches as a function of $k_7$ in the corresponding bifurcation diagrams. The columns are as follows: (1) variables; (2) absolute variation along the TNBC branch; (3) absolute variation along the HER2+ branch; (4) total absolute variation across the entire bifurcation diagram; (5-6) the percentage values, with column (4) representing 100%.

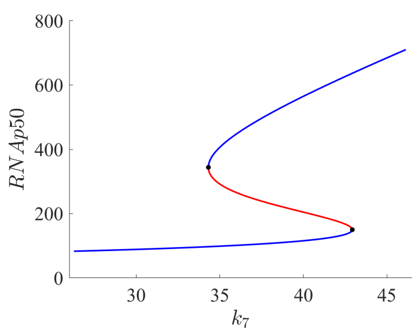


(a) $k_7 \times RNAp50$

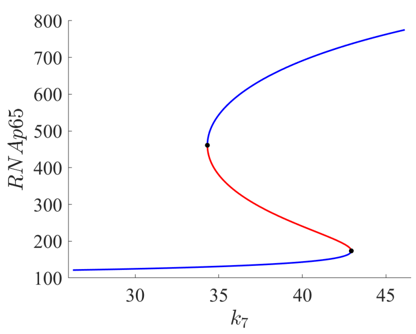


(b) $k_7 \times RNAp65$

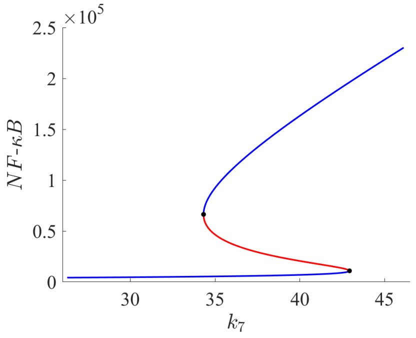


(c) $k_7 \times NF\text{-}\kappa B$

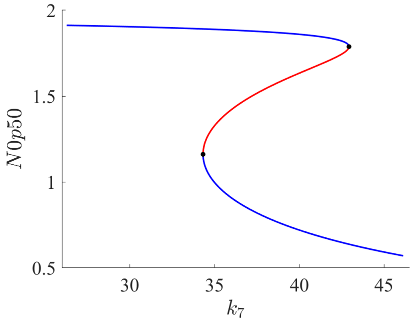


(d) $k_7 \times N0p50$

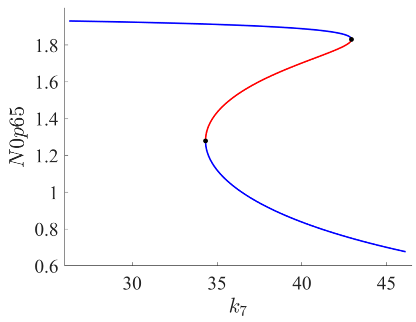


(e) $k_7 \times N0p65$

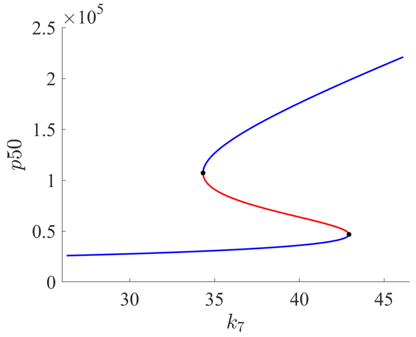


(f) $k_7 \times p50$

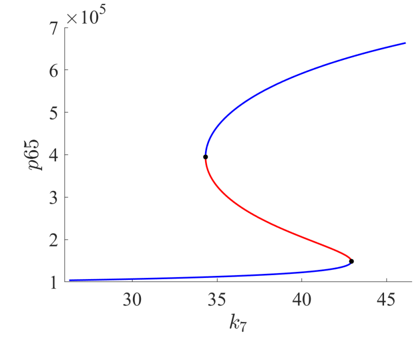


(g) $k_7 \times p65$

Figure S6: Bifurcation diagrams for all species with respect to the parameter $k_7$

| **Variables** | **TNBC** | **HER2+** | **Total** | **% TNBC** | **% HER2+** |
|---|---|---|---|---|---|
| $RNAp50$ | 169.54 | 41.63 | 452.80 | 37.44 | 9.19 |
| $RNAp65$ | 361.00 | 70.58 | 691.40 | 52.21 | 10.21 |
| $NF\text{-}\kappa B$ | 114905.70 | 6170.43 | 181597.28 | 63.28 | 3.40 |
| $N0p50$ | 0.46 | 0.11 | 1.24 | 37.44 | 9.19 |
| $N0p65$ | 0.69 | 0.14 | 1.33 | 52.21 | 10.21 |
| $p50$ | 52777.80 | 12959.13 | 140953.16 | 37.44 | 9.19 |
| $p65$ | 309049.57 | 60427.10 | 591904.43 | 52.21 | 10.21 |

Table S9: Variation of each species' copy number along the HER2+ and TNBC branches as a function of $k_8$ in the corresponding bifurcation diagrams. The columns are as follows: (1) variables; (2) absolute variation along the TNBC branch; (3) absolute variation along the HER2+ branch; (4) total absolute variation across the entire bifurcation diagram; (5-6) the percentage values, with column (4) representing 100%.

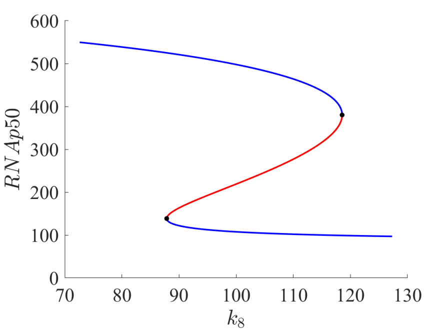


(a) $k_8 \times RNAp50$

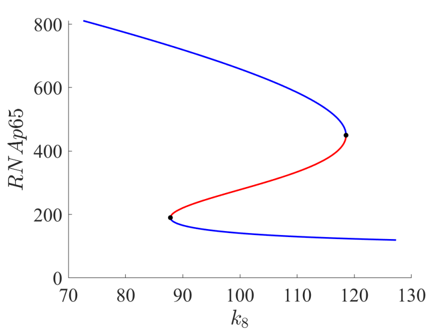


(b) $k_8 \times RNAp65$

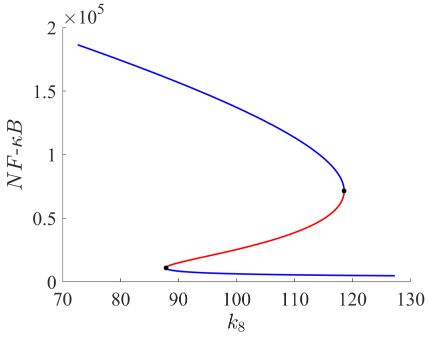


(c) $k_8 \times NF\text{-}\kappa B$

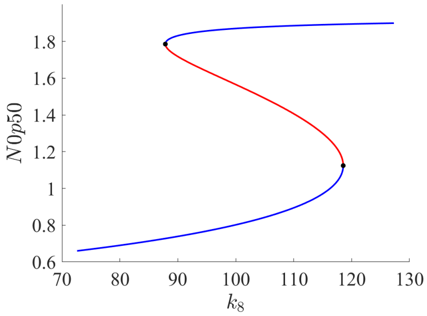


(d) $k_8 \times N0p50$

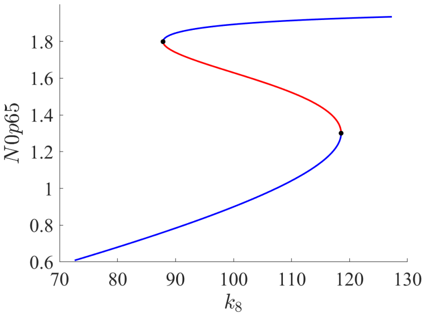


(e) $k_8 \times N0p65$

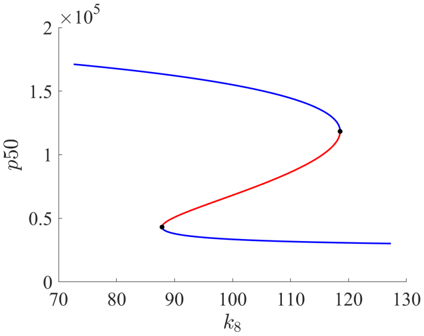


(f) $k_8 \times p50$

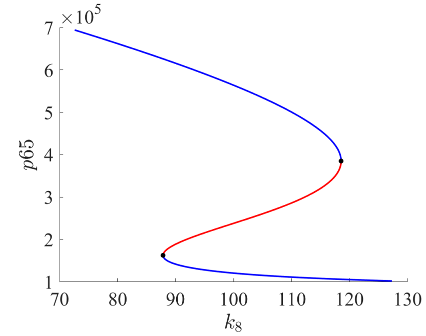


(g) $k_8 \times p65$

Figure S7: Bifurcation diagrams for all species with respect to the parameter $k_8$

| **Variables** | **TNBC** | **HER2+** | **Total** | **% TNBC** | **% HER2+** |
|---|---|---|---|---|---|
| $RNAp50$ | 166.44 | 44.18 | 452.25 | 36.80 | 9.77 |
| $RNAp65$ | 350.13 | 76.34 | 686.29 | 51.02 | 11.12 |
| $NF\text{-}\kappa B$ | 111368.55 | 6525.08 | 178414.77 | 62.42 | 3.66 |
| $N0p50$ | 0.46 | 0.12 | 1.24 | 36.80 | 9.77 |
| $N0p65$ | 0.67 | 0.15 | 1.32 | 51.02 | 11.12 |
| $p50$ | 51811.73 | 13754.41 | 140782.37 | 36.80 | 9.77 |
| $p65$ | 299746.08 | 65350.70 | 587524.54 | 51.02 | 11.12 |

Table S10: Variation of each species' copy number along the HER2+ and TNBC branches as a function of $k_9$ in the corresponding bifurcation diagrams. The columns are as follows: (1) variables; (2) absolute variation along the TNBC branch; (3) absolute variation along the HER2+ branch; (4) total absolute variation across the entire bifurcation diagram; (5-6) the percentage values, with column (4) representing 100%.

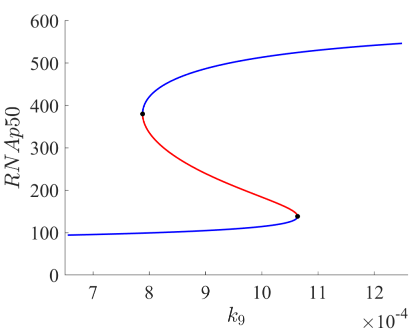


(a) $k_9 \times RNAp50$

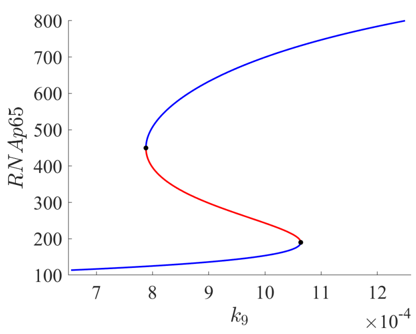


(b) $k_9 \times RNAp65$

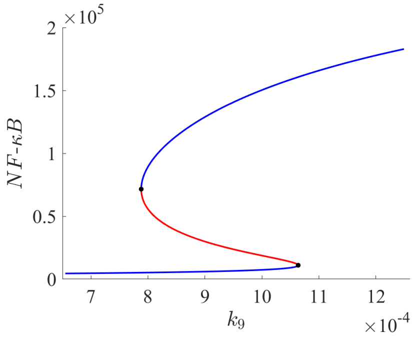


(c) $k_9 \times NF\text{-}\kappa B$

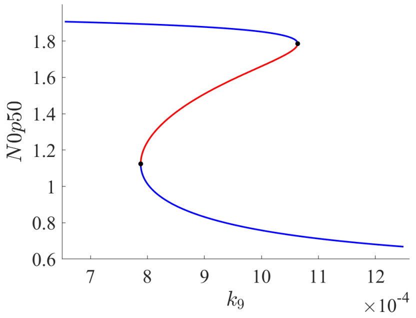


(d) $k_9 \times N0p50$

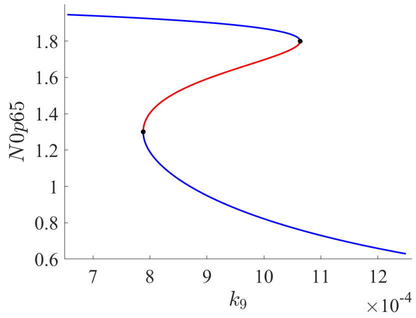


(e) $k_9 \times N0p65$

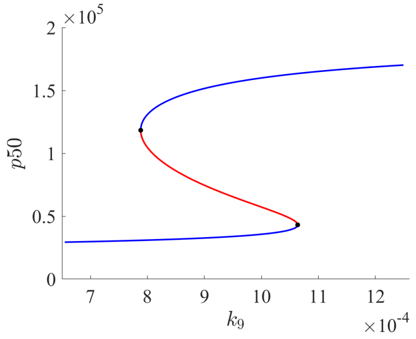


(f) $k_9 \times p50$

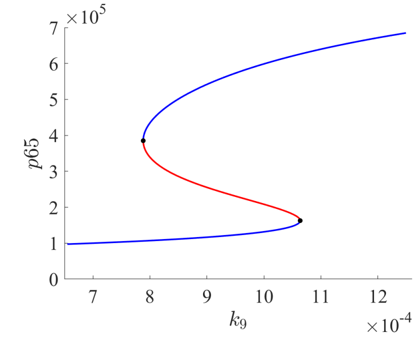


(g) $k_9 \times p65$

Figure S8: Bifurcation diagrams for all species with respect to the parameter $k_9$

| **Variables** | **TNBC** | **HER2+** | **Total** | **% TNBC** | **% HER2+** |
|---|---|---|---|---|---|
| $RNAp50$ | 236.89 | 42.33 | 508.28 | 46.61 | 8.33 |
| $RNAp65$ | 618.86 | 73.82 | 936.53 | 66.08 | 7.88 |
| $NF\text{-}\kappa B$ | 198733.50 | 6234.00 | 260238.73 | 76.37 | 2.40 |
| $N0p50$ | 0.65 | 0.12 | 1.39 | 46.61 | 8.33 |
| $N0p65$ | 0.67 | 0.09 | 1.31 | 50.80 | 7.13 |
| $p50$ | 73740.17 | 13178.13 | 158223.40 | 46.61 | 8.33 |
| $p65$ | 529803.17 | 63195.68 | 801753.04 | 66.08 | 7.88 |

Table S11: Variation of each species' copy number along the HER2+ and TNBC branches as a function of $k_{10}$ in the corresponding bifurcation diagrams. The columns are as follows: (1) variables; (2) absolute variation along the TNBC branch; (3) absolute variation along the HER2+ branch; (4) total absolute variation across the entire bifurcation diagram; (5-6) the percentage values, with column (4) representing 100%.

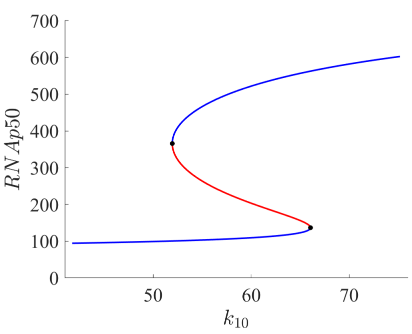


(a) $k_{10} \times RNAp50$

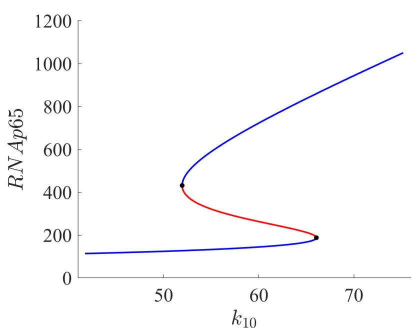


(b) $k_{10} \times RNAp65$

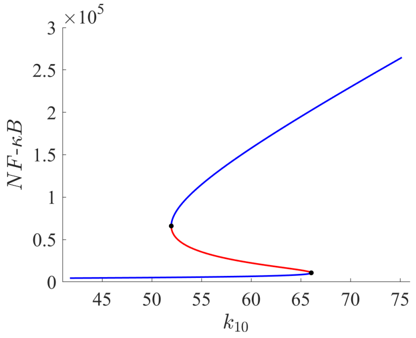


(c) $k_{10} \times NF\text{-}\kappa B$

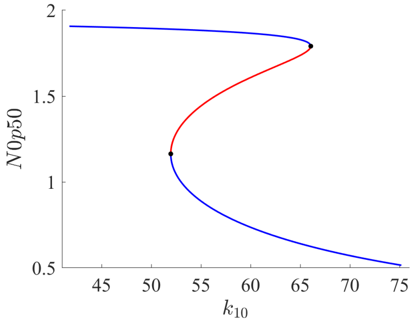


(d) $k_{10} \times N0p50$

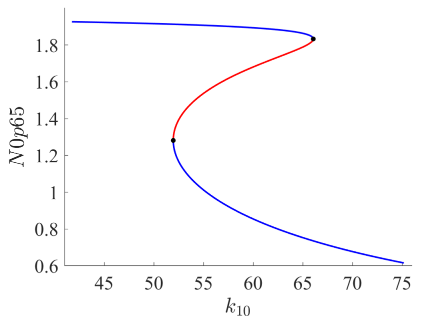


(e) $k_{10} \times N0p65$

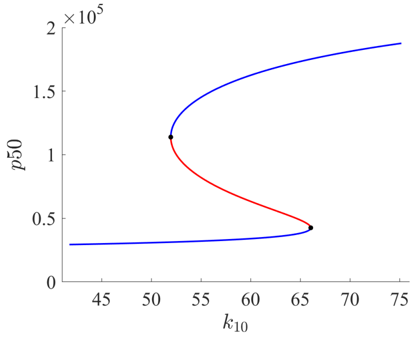


(f) $k_{10} \times p50$

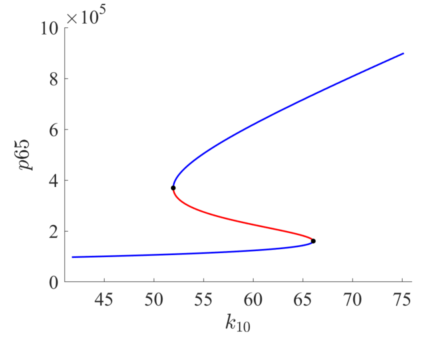


(g) $k_{10} \times p65$

Figure S9: Bifurcation diagrams for all species with respect to the parameter $k_{10}$

| **Variables** | **TNBC** | **HER2+** | **Total** | **% TNBC** | **% HER2+** |
|---|---|---|---|---|---|
| $RNAp50$ | 212.73 | 59.40 | 497.38 | 42.77 | 11.94 |
| $RNAp65$ | 310.17 | 68.41 | 662.54 | 46.81 | 10.33 |
| $NF\text{-}\kappa B$ | 163776.95 | 8759.72 | 228263.42 | 71.75 | 3.84 |
| $N0p50$ | 0.58 | 0.16 | 1.36 | 42.77 | 11.94 |
| $N0p65$ | 0.59 | 0.13 | 1.27 | 46.81 | 10.33 |
| $p50$ | 66220.58 | 18490.29 | 154830.23 | 42.77 | 11.94 |
| $p65$ | 265532.53 | 58566.08 | 567198.97 | 46.81 | 10.33 |

Table S12: Variation of each species' copy number along the HER2+ and TNBC branches as a function of $k_{11}$ in the corresponding bifurcation diagrams. The columns are as follows: (1) variables; (2) absolute variation along the TNBC branch; (3) absolute variation along the HER2+ branch; (4) total absolute variation across the entire bifurcation diagram; (5-6) the percentage values, with column (4) representing 100%.

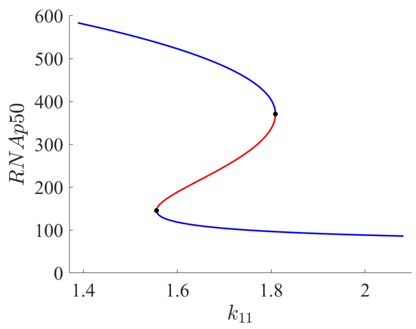


(a) $k_{11} \times RNAp50$

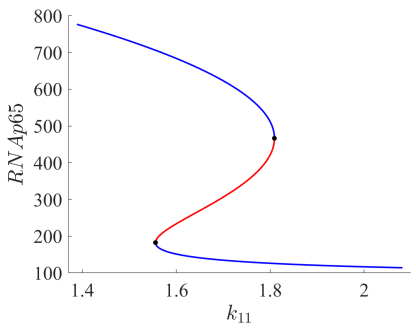


(b) $k_{11} \times RNAp65$

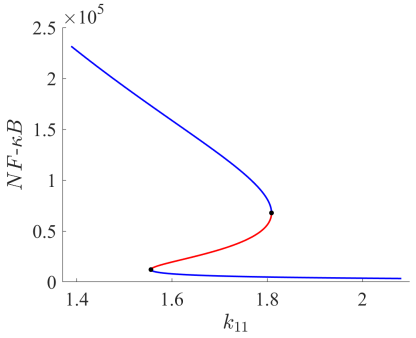


(c) $k_{11} \times NF\text{-}\kappa B$

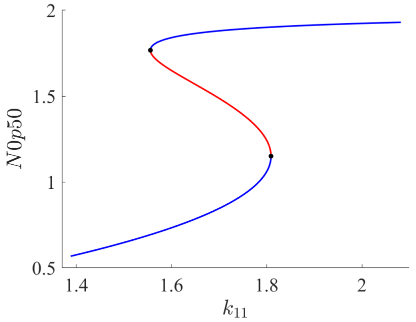


(d) $k_{11} \times N0p50$

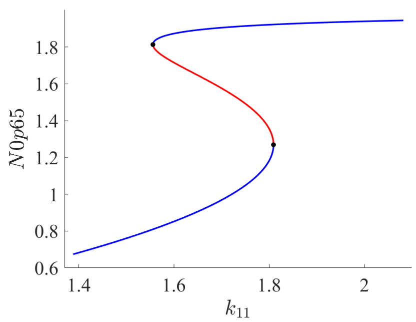


(e) $k_{11} \times N0p65$

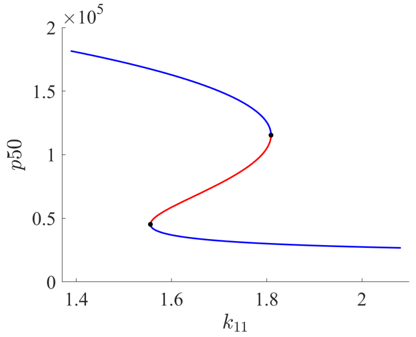


(f) $k_{11} \times p50$

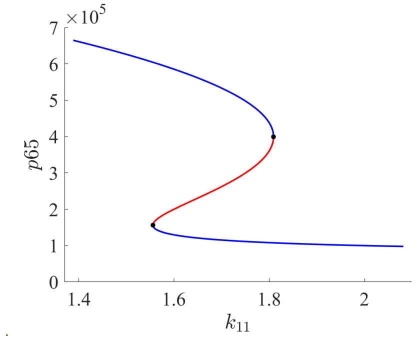


(g) $k_{11} \times p65$

Figure S10: Bifurcation diagrams for all species with respect to the parameter $k_{11}$

| **Variables** | **TNBC** | **HER2+** | **Total** | **% TNBC** | **% HER2+** |
|---|---|---|---|---|---|
| $RNAp50$ | 197.73 | 54.53 | 477.51 | 41.41 | 11.42 |
| $RNAp65$ | 286.75 | 62.90 | 633.62 | 45.26 | 9.93 |
| $NF\text{-}\kappa B$ | 141956.64 | 8097.95 | 205781.34 | 68.98 | 3.94 |
| $N0p50$ | 0.54 | 0.15 | 1.31 | 41.41 | 11.42 |
| $N0p65$ | 0.55 | 0.12 | 1.22 | 45.26 | 9.93 |
| $p50$ | 61550.30 | 16975.37 | 148645.04 | 41.41 | 11.42 |
| $p65$ | 245488.84 | 53847.29 | 542436.48 | 45.26 | 9.93 |

Table S13: Variation of each species' copy number along the HER2+ and TNBC branches as a function of $k_{12}$ in the corresponding bifurcation diagrams. The columns are as follows: (1) variables; (2) absolute variation along the TNBC branch; (3) absolute variation along the HER2+ branch; (4) total absolute variation across the entire bifurcation diagram; (5-6) the percentage values, with column (4) representing 100%.

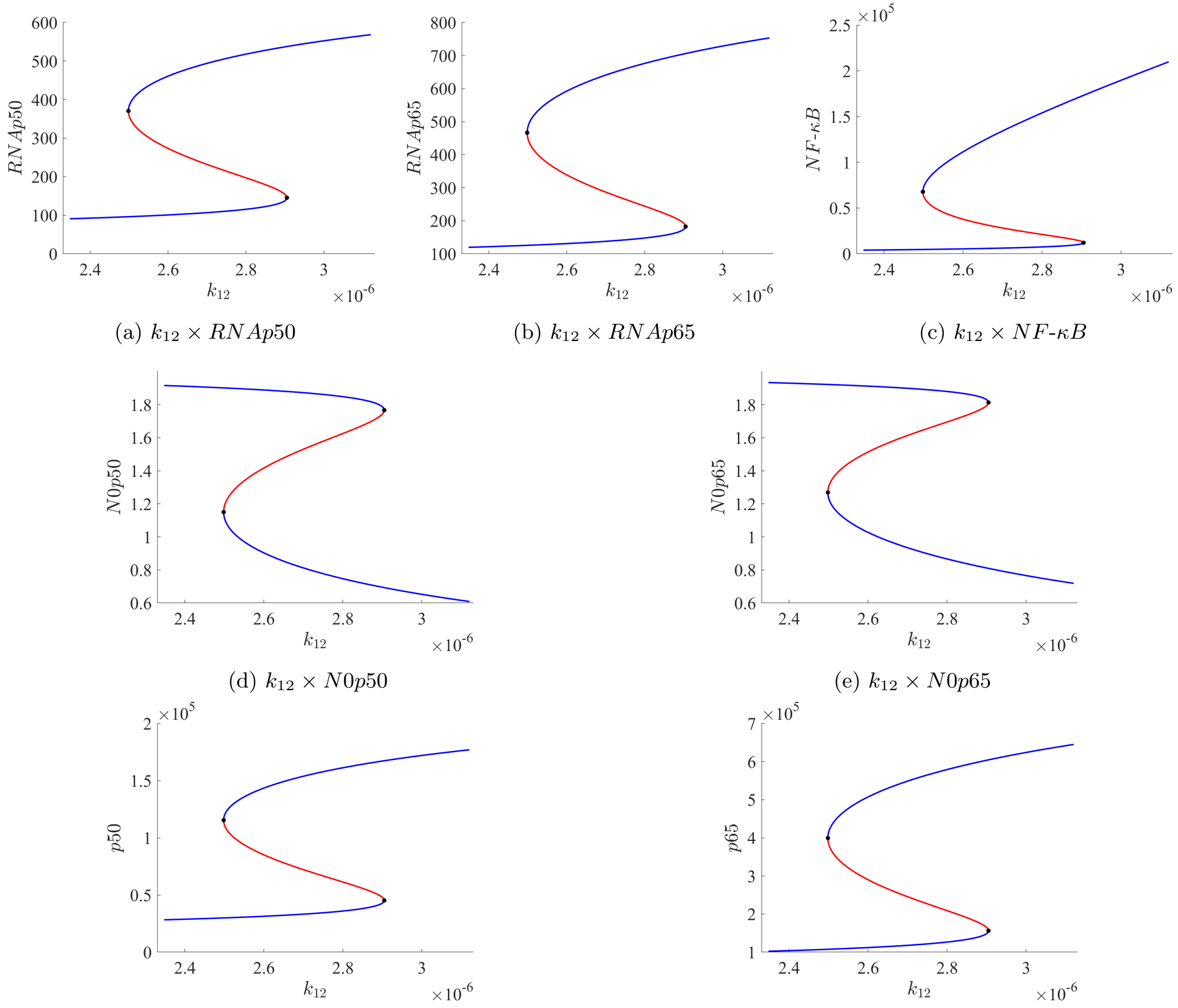


(a) $k_{12} \times RNAp50$ (b) $k_{12} \times RNAp65$ (c) $k_{12} \times NF\text{-}\kappa B$

(d) $k_{12} \times N0p50$ (e) $k_{12} \times N0p65$

(f) $k_{12} \times p50$ (g) $k_{12} \times p65$

Figure S11: Bifurcation diagrams for all species with respect to the parameter $k_{12}$

| **Variables** | **TNBC** | **HER2+** | **Total** | **% TNBC** | **% HER2+** |
|---|---|---|---|---|---|
| $RNAp50$ | 202.90 | 57.33 | 485.48 | 41.79 | 11.81 |
| $RNAp65$ | 294.80 | 66.07 | 644.83 | 45.72 | 10.25 |
| $NF\text{-}\kappa B$ | 149138.49 | 8479.66 | 213344.90 | 69.90 | 3.97 |
| $N0p50$ | 0.56 | 0.16 | 1.33 | 41.79 | 11.81 |
| $N0p65$ | 0.57 | 0.13 | 1.24 | 45.72 | 10.25 |
| $p50$ | 103894.22 | 26190.90 | 188946.30 | 54.99 | 13.86 |
| $p65$ | 252374.47 | 56562.82 | 552037.65 | 45.72 | 10.25 |

Table S14: Variation of each species' copy number along the HER2+ and TNBC branches as a function of $k_{13}$ in the corresponding bifurcation diagrams. The columns are as follows: (1) variables; (2) absolute variation along the TNBC branch; (3) absolute variation along the HER2+ branch; (4) total absolute variation across the entire bifurcation diagram; (5-6) the percentage values, with column (4) representing 100%.

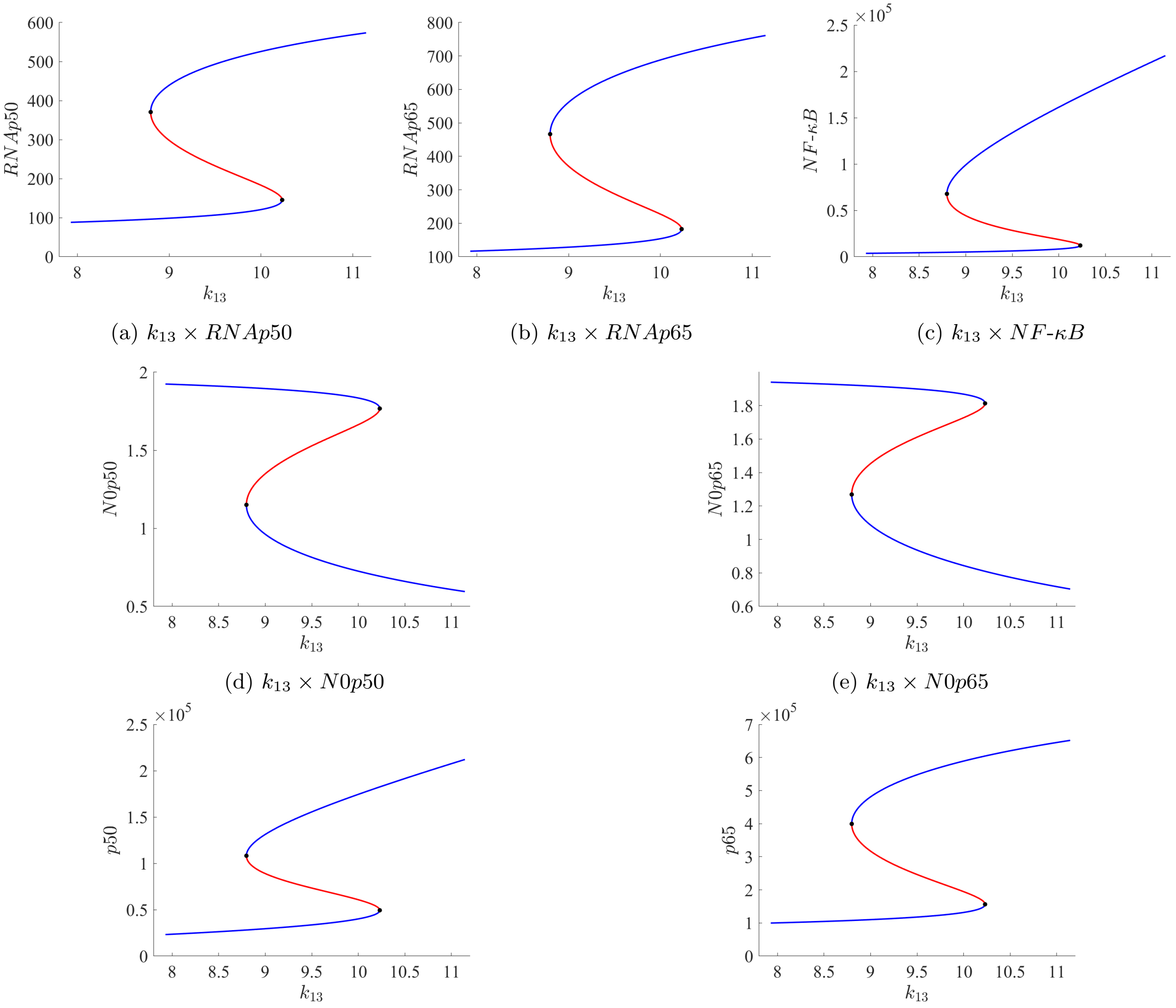


(a) $k_{13} \times RNAp50$ (b) $k_{13} \times RNAp65$ (c) $k_{13} \times NF\text{-}\kappa B$

(d) $k_{13} \times N0p50$ (e) $k_{13} \times N0p65$

(f) $k_{13} \times p50$ (g) $k_{13} \times p65$

Figure S12: Bifurcation diagrams for all species with respect to the parameter $k_{13}$

| **Variables** | **TNBC** | **HER2+** | **Total** | **% TNBC** | **% HER2+** |
|---|---|---|---|---|---|
| $RNAp50$ | 190.75 | 53.49 | 469.49 | 40.63 | 11.39 |
| $RNAp65$ | 275.95 | 61.72 | 621.63 | 44.39 | 9.93 |
| $NF\text{-}\kappa B$ | 132781.28 | 7954.93 | 196462.95 | 67.59 | 4.05 |
| $N0p50$ | 0.52 | 0.15 | 1.29 | 40.63 | 11.39 |
| $N0p65$ | 0.53 | 0.12 | 1.19 | 44.39 | 9.93 |
| $p50$ | 59378.32 | 16650.72 | 146148.41 | 40.63 | 11.39 |
| $p65$ | 356829.63 | 77983.53 | 638985.45 | 55.84 | 12.20 |

Table S15: Variation of each species' copy number along the HER2+ and TNBC branches as a function of $k_{14}$ in the corresponding bifurcation diagrams. The columns are as follows: (1) variables; (2) absolute variation along the TNBC branch; (3) absolute variation along the HER2+ branch; (4) total absolute variation across the entire bifurcation diagram; (5-6) the percentage values, with column (4) representing 100%.

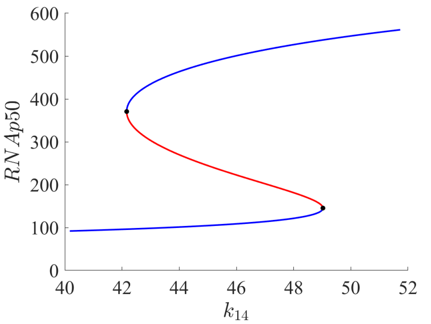


(a) $k_{14} \times RNAp50$

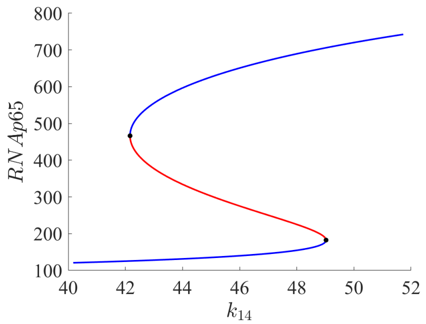


(b) $k_{14} \times RNAp65$

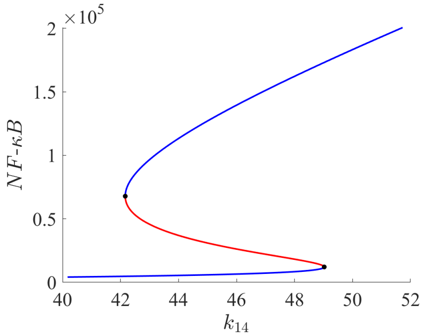


(c) $k_{14} \times NF\text{-}\kappa B$

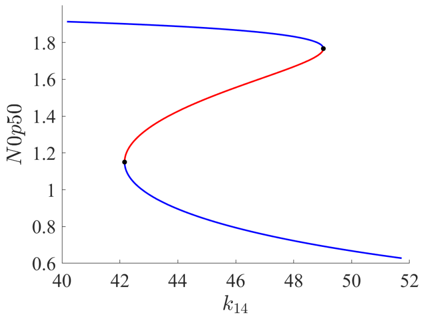


(d) $k_{14} \times N0p50$

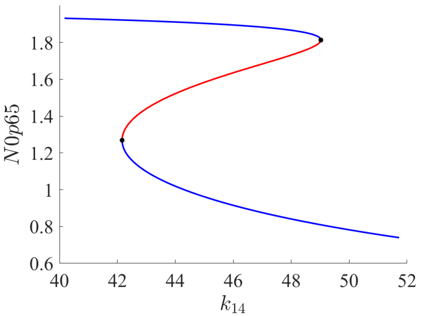


(e) $k_{14} \times N0p65$

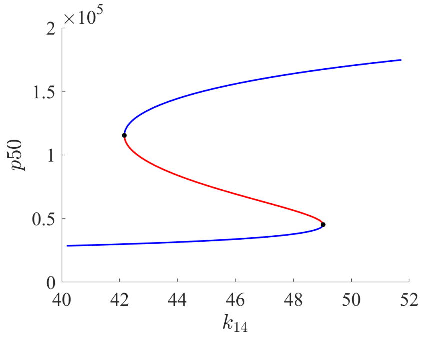


(f) $k_{14} \times p50$

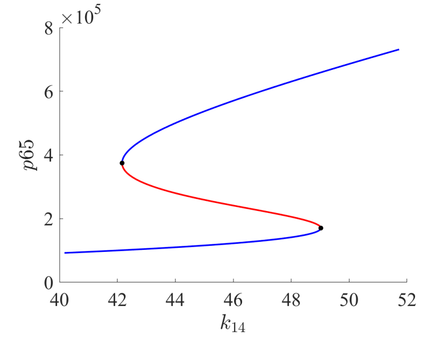


(g) $k_{14} \times p65$

Figure S13: Bifurcation diagrams for all species with respect to the parameter $k_{14}$

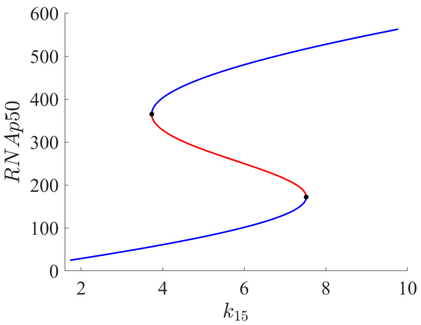


(a) $k_{15} \times RNAp50$

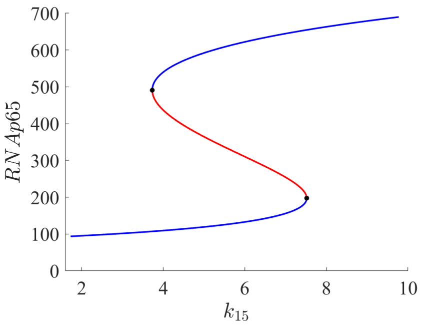


(b) $k_{15} \times RNAp65$

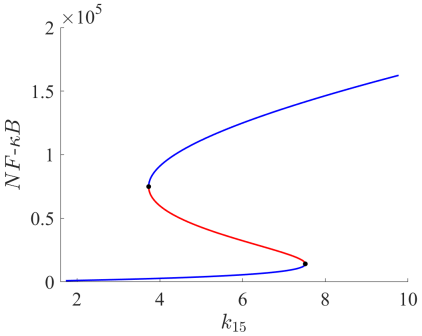


(c) $k_{15} \times NF\text{-}\kappa B$

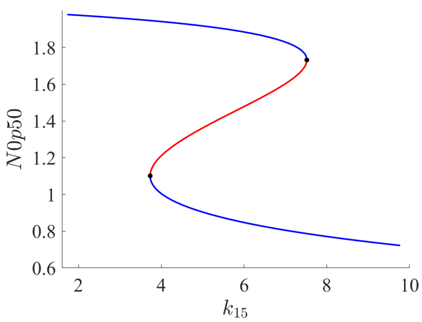


(d) $k_{15} \times N0p50$

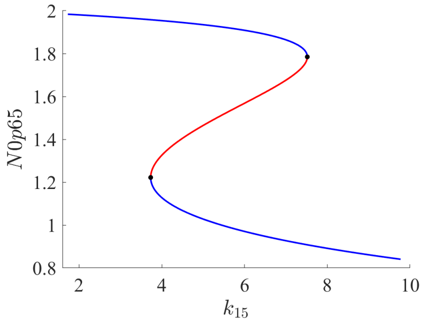


(e) $k_{15} \times N0p65$

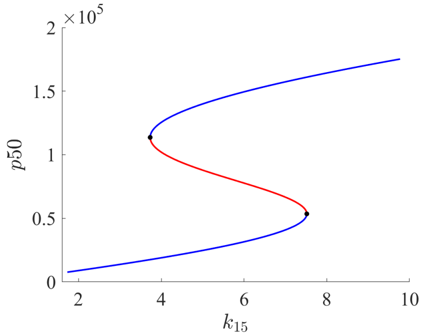


(f) $k_{15} \times p50$

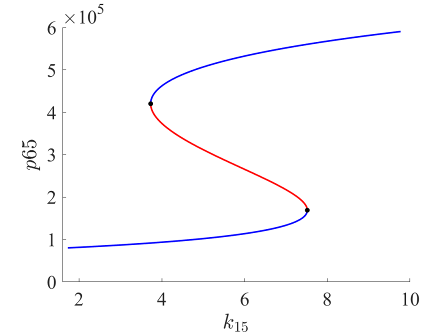


(g) $k_{15} \times p65$

Figure S14: Bifurcation diagrams for all species with respect to the parameter $k_{15}$

| Variables | TNBC | HER2+ | Total | % TNBC | % HER2+ |
|---|---|---|---|---|---|
| $RNAp50$ | 197.85 | 147.22 | 538.22 | 36.76 | 27.35 |
| $RNAp65$ | 198.91 | 103.74 | 596.04 | 33.37 | 17.41 |
| $NF\text{-}\kappa B$ | 87454.53 | 13233.05 | 161439.50 | 54.17 | 8.20 |
| $N0p50$ | 0.38 | 0.25 | 1.26 | 30.14 | 19.65 |
| $N0p65$ | 0.38 | 0.20 | 1.14 | 33.37 | 17.41 |
| $p50$ | 61589.40 | 45828.93 | 167542.32 | 36.76 | 27.35 |
| $p65$ | 170282.47 | 88813.00 | 510269.31 | 33.37 | 17.41 |

Table S16: Variation of each species' copy number along the HER2+ and TNBC branches as a function of $k_{15}$ in the corresponding bifurcation diagrams. The columns are as follows: (1) variables; (2) absolute variation along the TNBC branch; (3) absolute variation along the HER2+ branch; (4) total absolute variation across the entire bifurcation diagram; (5-6) the percentage values, with column (4) representing 100%.Variation of each species' copy number along the HER2+ and TNBC branches as a function of $k_1$ in the corresponding bifurcation diagrams. The columns are as follows: (1) variables; (2) absolute variation along the TNBC branch; (3) absolute variation along the HER2+ branch; (4) total absolute variation across the entire bifurcation diagram; (5-6) the percentage values, with column (4) representing 100%.

| Variables | TNBC | HER2+ | Total | % TNBC | % HER2+ |
|---|---|---|---|---|---|
| $RNAp50$ | 128.75 | 89.80 | 451.11 | 28.54 | 19.91 |
| $RNAp65$ | 250.11 | 181.13 | 680.48 | 36.75 | 26.62 |
| $NF\text{-}\kappa B$ | 79196.38 | 13150.61 | 153795.05 | 51.49 | 8.55 |
| $N0p50$ | 0.35 | 0.25 | 1.24 | 28.54 | 19.91 |
| $N0p65$ | 0.35 | 0.20 | 1.12 | 31.62 | 17.68 |
| $p50$ | 40077.98 | 27954.01 | 140425.88 | 28.54 | 19.91 |
| $p65$ | 214115.25 | 155062.26 | 582556.03 | 36.75 | 26.62 |

Table S17: Variation of each species' copy number along the HER2+ and TNBC branches as a function of $k_{16}$ in the corresponding bifurcation diagrams. The columns are as follows: (1) variables; (2) absolute variation along the TNBC branch; (3) absolute variation along the HER2+ branch; (4) total absolute variation across the entire bifurcation diagram; (5-6) the percentage values, with column (4) representing 100%.

| $\mathbf{k_3}$ | RNAp50 | RNAp65 | NF-κB | N0p50 | N0p65 | p50 | p65 |
|---|---|---|---|---|---|---|---|
| 0.054321555749 | 99.55 | 129.48 | 5211.47 | 1.89 | 1.92 | 30989.97 | 107145.85 |
| | 286.70 | 355.80 | 41239.94 | 1.38 | 1.48 | 89246.62 | 294416.94 |
| | 449.33 | 575.97 | 104622.88 | 0.93 | 1.059 | 139871.98 | 476576.32 |
| 0.0511933259 | 109.12 | 140.43 | 6573.79 | 1.87 | 1.90 | 33968.83 | 123302.52 |
| | 220.47 | 272.65 | 25786.79 | 1.56 | 1.64 | 68630.66 | 239395.36 |
| | 502.36 | 653.11 | 140750.57 | 0.79 | 0.91 | 156381.74 | 573456.70 |

Table S18: Stationary states for different values of $k_3$. For each $k_3$, the first row corresponds to the HER2+ subtype, the second to the unstable state, and the third to the TNBC subtype.

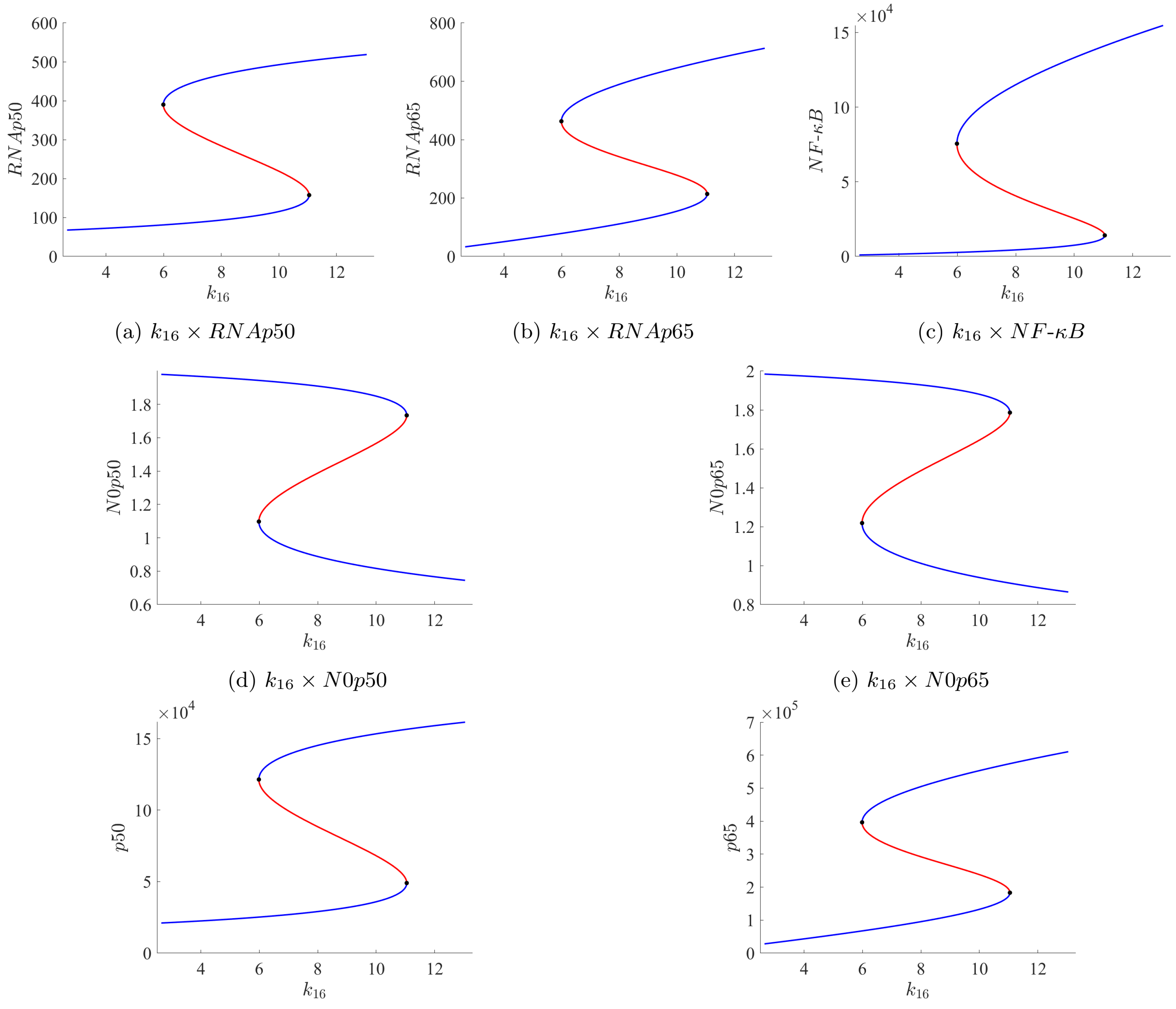


(a) $k_{16} \times RNAp50$ (b) $k_{16} \times RNAp65$ (c) $k_{16} \times NF\text{-}\kappa B$

(d) $k_{16} \times N0p50$ (e) $k_{16} \times N0p65$

(f) $k_{16} \times p50$ (g) $k_{16} \times p65$

Figure S15: Bifurcation diagrams for all species with respect to the parameter $k_{16}$

| $\mathbf{k_{12}}$ | **RNAp50** | **RNAp65** | **NF-$\kappa$B** | **N0p50** | **N0p65** | **p50** | **p65** |
|---|---|---|---|---|---|---|---|
| $2.725683 \times 10^{-6}$ | 108.67 | 139.91 | 6508.46 | 1.87 | 1.90 | 33827.85 | 119775.32 |
| | 222.50 | 275.15 | 26207.81 | 1.56 | 1.64 | 69263.86 | 235552.49 |
| | 500.91 | 650.95 | 139582.89 | 0.79 | 0.92 | 155928.74 | 557275.12 |
| $2.574339 \times 10^{-6}$ | 99.56 | 129.49 | 5212.53 | 1.89 | 1.92 | 30992.33 | 110858.22 |
| | 286.61 | 355.69 | 41216.46 | 1.38 | 1.48 | 89218.92 | 304499.76 |
| | 449.41 | 576.05 | 104667.48 | 0.93 | 1.059 | 139896.11 | 493150.44 |

Table S19: Stationary states for different values of $k_{12}$. For each $k_{12}$, the first row corresponds to the HER2+ subtype, the second to the unstable state, and the third to the TNBC subtype.

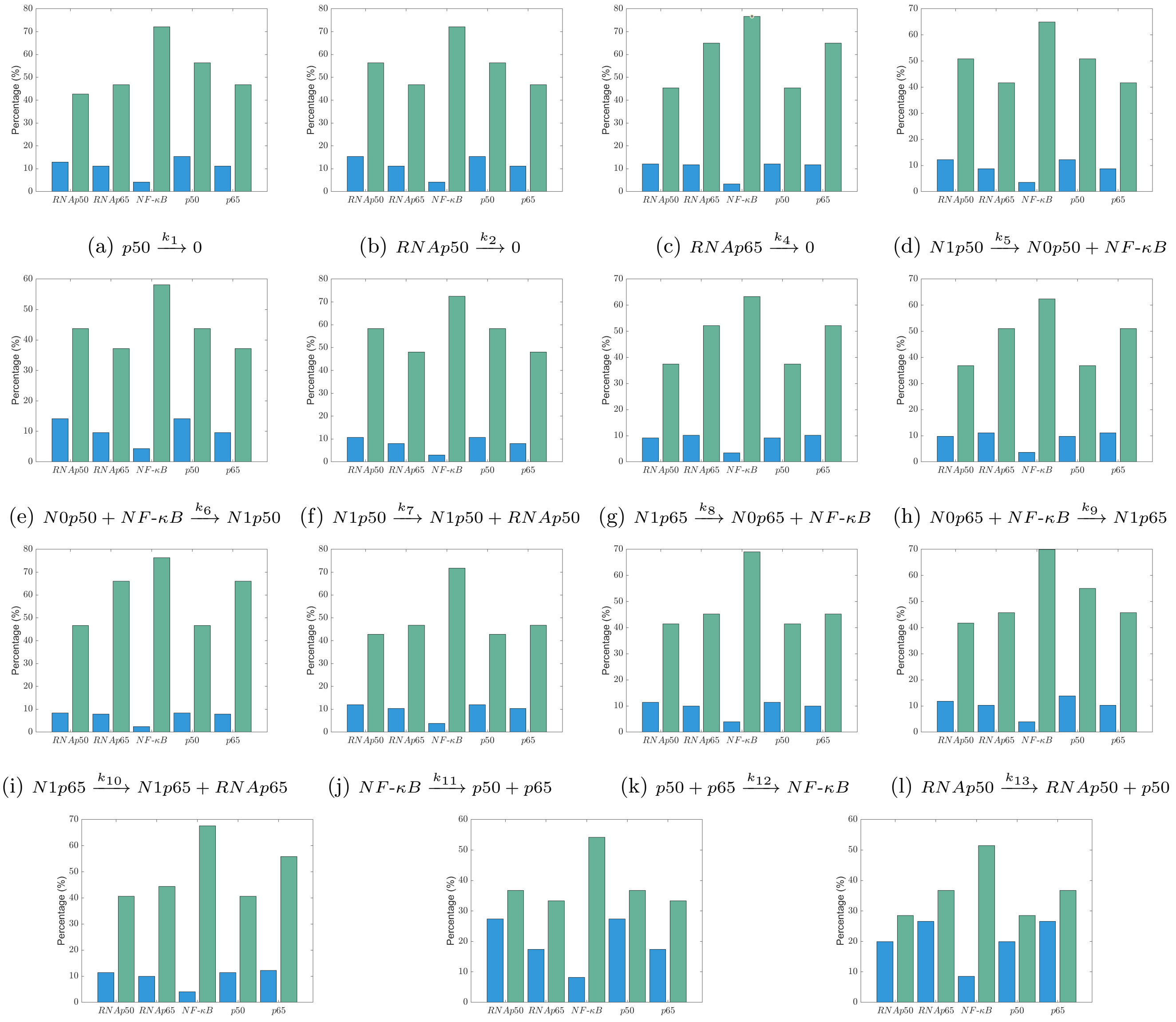


(a) $p50 \xrightarrow{k_1} 0$
(b) $RNAp50 \xrightarrow{k_2} 0$
(c) $RNAp65 \xrightarrow{k_4} 0$
(d) $N1p50 \xrightarrow{k_5} N0p50 + NF\text{-}\kappa B$
(e) $N0p50 + NF\text{-}\kappa B \xrightarrow{k_6} N1p50$
(f) $N1p50 \xrightarrow{k_7} N1p50 + RNAp50$
(g) $N1p65 \xrightarrow{k_8} N0p65 + NF\text{-}\kappa B$
(h) $N0p65 + NF\text{-}\kappa B \xrightarrow{k_9} N1p65$
(i) $N1p65 \xrightarrow{k_{10}} N1p65 + RNAp65$
(j) $NF\text{-}\kappa B \xrightarrow{k_{11}} p50 + p65$
(k) $p50 + p65 \xrightarrow{k_{12}} NF\text{-}\kappa B$
(l) $RNAp50 \xrightarrow{k_{13}} RNAp50 + p50$
(m) $RNAp65 \xrightarrow{k_{14}} RNAp65 + p65$
(n) $0 \xrightarrow{k_{15}} RNAp50$
(o) $0 \xrightarrow{k_{16}} RNAp65$

Figure S16: Sensitivity analysis of species copy-number variations with respect to each kinetic parameter $k_i$ ($i = 1, \ldots, 16$). Each panel corresponds to a single reaction in the regulatory network. Blue bars represent variations along the HER2+ branch, and green bars represent variations along the TNBC branch.

### Bifurcation diagrams with two free parameters

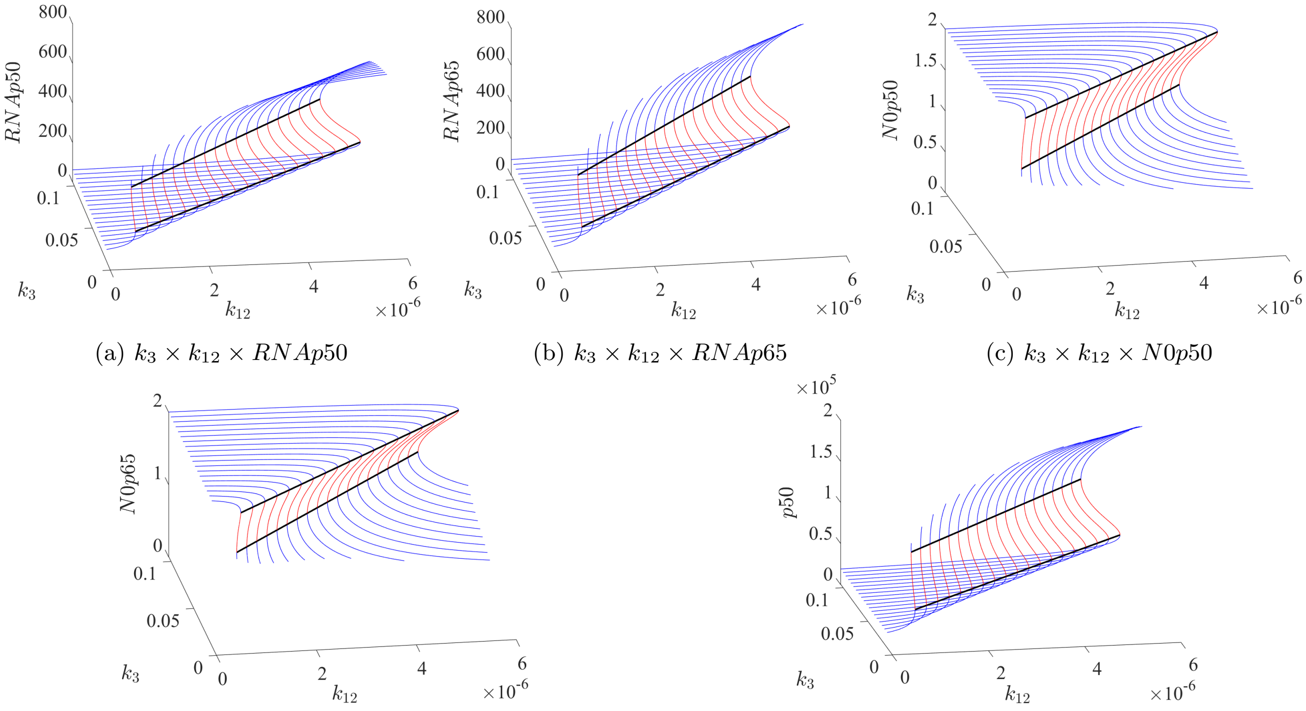


(a) $k_3 \times k_{12} \times RNAp50$ (b) $k_3 \times k_{12} \times RNAp65$ (c) $k_3 \times k_{12} \times N0p50$

(d) $k_3 \times k_{12} \times N0p65$ (e) $k_3 \times k_{12} \times p50$

Figure S17: Three-dimensional bifurcation diagrams with respect to parameters $k_3$ and $k_{12}$, illustrating the stationary-state structure of RNAp50, RNAp65, N0p50, N0p65, and p50 (panels a–e).
